# Engineered chemotaxis core signaling units indicate a constrained kinase-off state

**DOI:** 10.1101/2020.03.27.011866

**Authors:** Alise R. Muok, Teck Khiang Chua, Madhur Srivastava, Wen Yang, Zach Maschmann, Petr P. Borbat, Jenna Chong, Sheng Zhang, Jack H. Freed, Ariane Briegel, Brian R. Crane

## Abstract

Bacterial chemoreceptors, the CheA histidine kinase, and the coupling protein CheW comprise transmembrane molecular arrays with remarkable sensing properties. An unanswered question concerns how receptors turn off CheA kinase activity. Chemoreceptor cytoplasmic regions engineered to assume a trimer-of-receptor-dimers configuration form well-defined complexes with CheA and CheW and promote a kinase-off state. These mimics of core signaling units were assembled to homogeneity and investigated by site-directed spin-labeling with pulse-dipolar ESR spectroscopy (PDS), small-angle x-ray scattering, targeted protein cross-linking, and cryo-electron microscopy. The kinase-off state is especially stable, has relatively low domain mobility and associates the histidine substrate domain P1 and docking domain P2 with the kinase core. Distances measured between spin-labeled ADP molecules bound to the P4 kinase domain provide evidence for a “dipped conformation” that has been previously proposed from molecular dynamics simulations. Taken together, the data provide an experimentally restrained model for the inhibited state of the core-signaling unit and suggest that chemoreceptors indirectly sequester the kinase and substrate domains to limit histidine autophosphorylation.

## Introduction

Many bacteria employ a complex transmembrane sensory apparatus to modulate their motility in response to the chemical environment (reviewed in (Falke, JJ, et al., 2014; Hazelbauer, GL, et al., 2010; Muok, AR, et al., 2019; Parkinson, JS, et al., 2015). Extensively studied in *Escherichia coli* (*Ec*), bacterial chemotaxis relies on transmembrane chemoreceptors to transduce extracellular signals into intracellular phosphorylation events (Falke, JJ, et al., 2014; Hazelbauer, GL, et al., 2010; Muok, AR, et al., 2019; Parkinson, JS, et al., 2015). The receptors, also known as methyl-accepting chemotaxis proteins (MCPs), organize into a trimer-of-dimers (TOD) that further assemble with the dimeric histidine kinase CheA and coupling protein CheW at their membrane-distal tips to produce an extended molecular lattice capable of highly sensitive, cooperative responses (Mello, BA, et al., 2003; Pinas, GE, et al., 2016; Sourjik, V, et al., 2002). MCPs regulate CheA autophoshorylation rates in response to binding signals from external ligands. Phosphoryl transfer from CheA to CheY activates CheY to modulate flagellar rotation sense. MCPs also undergo methylation and demethylation on specific glutamate residues by the methyl transferase CheR and the methyl esterase CheB, respectively. In *E. coli*, MCP methylation activates CheA and thereby opposes the action of attractant binding, which inhibits the kinase. The methylation system thus adapts the array output to the evolving ligand environment.

Despite considerable advances, it remains unclear how CheA is regulated by receptors (Falke, JJ, et al., 2014; Hazelbauer, GL, et al., 2010; Muok, AR, et al., 2019; Parkinson, JS, et al., 2015). CheA is composed of five domains (P1 - P5), each with a distinct function: P1 contains the phosphate-accepting histidine residue, P2 docks CheY, P3 dimerizes the subunits, P4 binds ATP as a member of the GHL family of ATPases, and P5 interacts with CheW to form molecular rings integral to the array architecture. The CheA P1 domain contains the substrate histidine residue (His48 in *Ec* CheA and His45 in *Thermotoga maritima* (*Tm*) CheA) and consists of 5 *α−*helices (A-E), with A-D composing a four-helix bundle and helix E connected to the bundle by a flexible linker (Quezada, CM, et al., 2004; Zhou, H, et al., 1995). The CheA domains are quite flexible in the free kinase, with the P1 and P2 modules sampling considerable conformational space (Bhatnagar, J, et al., 2010; Gloor, SL, et al., 2009; Greenswag, AR, et al., 2015). However, incorporation of CheA into the receptor arrays constrains its conformation (Briegel, A, et al., 2012; Cassidy, CK, et al., 2015; Cassidy, CK, et al., 2020; Yang, W, et al., 2019).

X-ray crystallography of composite domains and complexes, cryo-electron-tomography (cryo-ET) of native and ex-vivo assembled arrays, biochemistry, spectroscopic approaches, genetics and MD simulations have been combined to define the architecture and sensing behavior of the arrays (reviewed in (Falke, JJ, et al., 2014; Muok, AR, et al., 2019; Parkinson, JS, et al., 2015)): CheA sits at the interface of two TODs, with the helical P3 domain projecting between them toward the membrane. Paralogous P5 and CheW interact through conserved surfaces at the ends of their *β*-barrels to form rings composed from two types of interfaces: interface 1, which is proximal to P3 and involves P5 subdomain 1 and CheW subdomain 2; and interface 2, which is distal from P3 and involves P5 subdomain 2 and CheW subdomain 1. Dimeric MCPs interact in grooves at the center of CheW and P5 on the external surfaces of the rings. CheW-only rings may form on the 6-fold symmetry axis of the hexagonal lattice (Cassidy, CK, et al., 2015). The P4 kinase domains suspend below the receptor tips and align underneath the P5 domains. Long, flexible linkers connect P1 and P2 to each other (L1) and P2 to P3 (L2), whereas linkers between P3 and P4 (L3), and P4 and P5 (L4), play an important role in kinase activity and regulation (Ding, XY, et al., 2018; Wang, XQ, et al., 2014; Wang, XQ, et al., 2012). Positions of the P1 and P2 domains are not formally known, although cryo-ET suggests that they reside below the P4 domains of deactivated kinase (Briegel, A, et al., 2013; Yang, W, et al., 2019). However, the P1 and P2 positions are uncertain owing to limiting resolution in the cryo-ET reconstructions and/or domain mobility. The P4 domains are also not fully discernible in the tomograms of either *in vivo* or reconstituted arrays, but appear more ordered with the inactive kinase (Briegel, A, et al., 2013; Briegel, A, et al., 2012; Cassidy, CK, et al., 2015; Liu, J, et al., 2012; Yang, W, et al., 2019). Recent MD-assisted modeling against 12 Å resolution cryo-ET data has better defined two predominant conformations of the P4 domain below the P5-CheW layer and suggests that kinase regulation may involve transitions between these conformations (Cassidy, CK, et al., 2020).

The ability of receptors to alter *Ec* CheA phosphotransfer rates by several hundred-fold suggests that the kinase assumes substantially different structural and dynamical states associated with these activity changes (Falke, JJ, et al., 2014; Hazelbauer, GL, et al., 2010; Muok, AR, et al., 2019; Parkinson, JS, et al., 2015). Activity assays with P1 supplied as a separated substrate to the core kinase (domains P3P4P5) in complex with receptors indicate that activation primarily derives from increases in k_cat_ and not changes to the P1 Michaelis constant (K_M_) (Mello, BA, et al., 2018; Pan, W, et al., 2017). Furthermore, both ATP-binding and autophosphorylation (which includes both P1 binding and phosphate transfer) are steps potentially regulated by receptors (Mello, BA, et al., 2018). Interestingly, removal of one of the two P1 domains from a CheA dimer increases autophosphorylation activity, suggesting that the P1 domains may interfere with each other (Greenswag, AR, et al., 2015; Levit, M, et al., 1996). Molecular dynamic simulations of array models based on cryo-ET data indicate that the P4 domains “dip” toward each other from a position where the P4 ATP-binding pocket and flexible ATP “lid” reside near the P5-CheW layer (Cassidy, CK, et al., 2015; Cassidy, CK, et al., 2020). It follows that changes to kinase activity level relate to P4 location or motion.

Despite the extensive coupling of components within the natural arrays, a core-signaling unit comprised of two TODs, one dimeric CheA and two CheW proteins embodies considerable functionality (Frank, V, et al., 2016; Li, M, et al., 2014; Li, MS, et al., 2011; Pinas, GE, et al., 2016). In efforts to reconstitute a homogenous signaling particle of these components we previously developed a receptor engineering approach, wherein single chain versions of cytoplasmic receptor signaling domain “dimers” were trimerized by the foldon domain from bacteriophage T4 fibritin (Greenswag, AR, et al., 2015). These so-called receptor foldons produce trimeric species that are capable of modulating CheA activity (Greenswag, AR, et al., 2015). We now extend the foldon approach to reconstitute core signaling units in a highly inhibited state and characterize this species by a variety of biophysical methods, including SAXS, interface analysis by cross-linking and mass-spectrometry, and pulse-dipolar ESR spectroscopy (PDS). The results show that inhibited CheA exhibits increased interactions among the P1, P2 and P4 domains and relatively well-ordered L1 and L2 connecting loops. Furthermore, we provide direct evidence for the P4 “dipped conformation” observed in MD simulations (Cassidy, CK, et al., 2015) and compose models of the foldon-associated signaling particles that capture this property.

## Results

### Incorporation of TOD receptor foldons into ternary complexes

Different types of single-chain receptors have been employed to modulate CheA activity (Greenswag, AR, et al., 2015; Mowery, P, et al., 2015). Soluble chimeric receptors fragments containing a non-native trimerization motif pre-form a TOD arrangement and associate with CheA and CheW (Greenswag, AR, et al., 2015). The variants consist of a fused dyad of receptor cytoplasmic domains (each 71 residues) from either the *Tm* MCP Tm14 or *Ec* MCP Tar linked at the C-terminus to a “fold-on” trimerization motif from the T4 phage protein fibritin (Figure 1A,B). The receptor segment includes the ∼36 residue highly conserved protein interaction region (PIR) that interacts with CheW and P5 and mediates the TOD trimer contact. Despite deriving from distantly related bacteria, the Tm14 and Tar foldons share 44% identity and 67% sequence homology over their 71 residue sequences, including 72% identity and 86% homology over the 36 residue PIR that binds to CheA and CheW. Previous small-angle x-ray scattering (SAXS) data and spin-labeling studies with the Tar foldon revealed a compact, globular shape with associated tips (Greenswag, AR, et al., 2015). Multi-angle light scattering (MALS) experiments with the purified foldons confirm trimerization in solution (Figure 1 – figure supplement 1A,C) and stable interactions with CheA and CheW (Figure 1 – figure supplement 1B,D).

**Figure 1.**
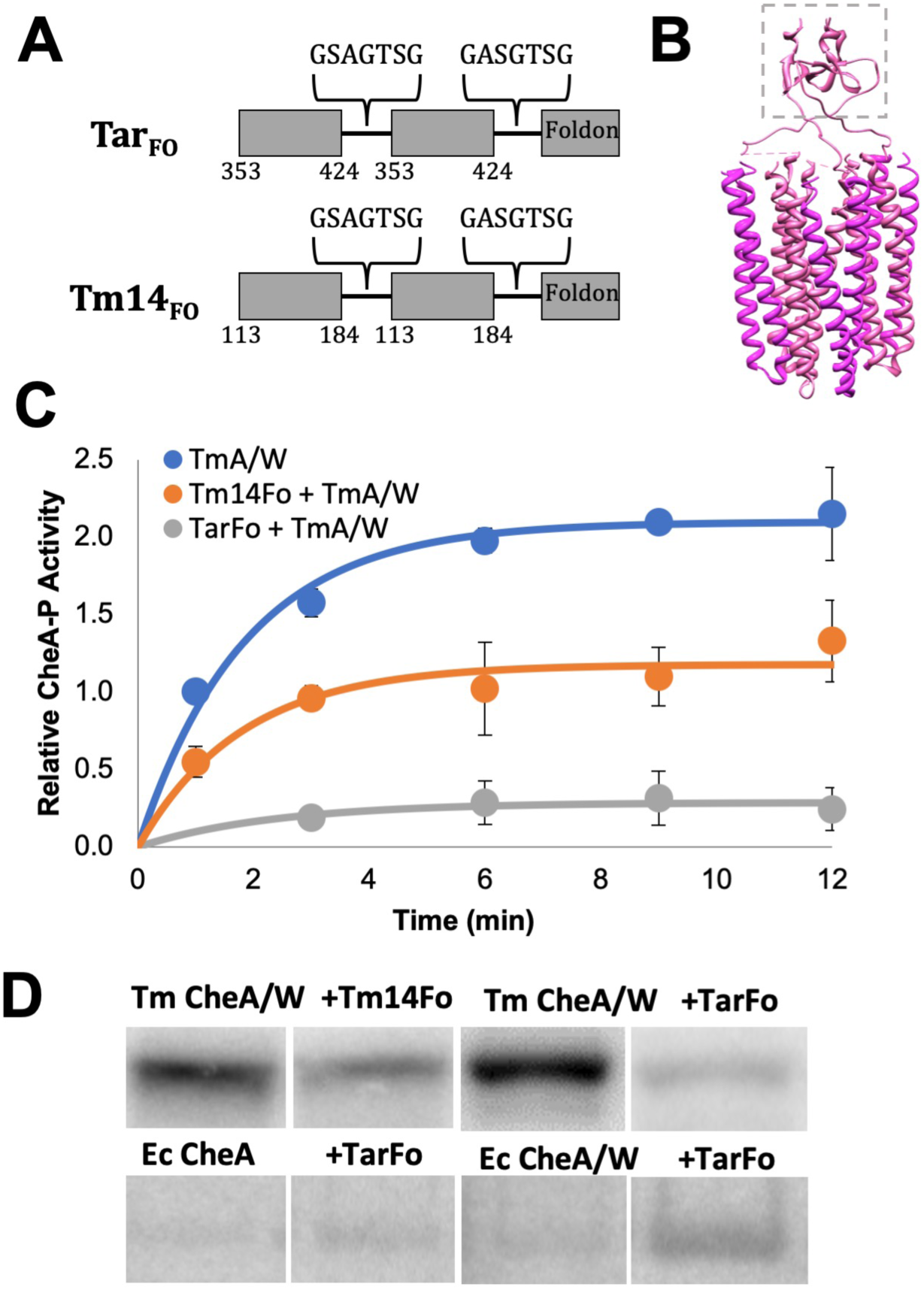
Construction and characterization of the Tar and Tm14 foldons. (A) The foldons were engineered to produce a pre-formed trimer-of-(single-chain)-dimer arrangement. Receptor regions surrounding the PIRs were fused to form a single-chained dimer that is capped with an N-terminal trimerization motif from T4-phage fibritin. (B) Model of the receptor foldons with the fibritin trimerization motif depicted in a dashed box. (C) Both the Tm14 foldon and Tar foldon deactivate *Tm* CheA autophosphorylation. Curves were fit to a = a0(1-exp(kt)), a0*k represents the relative initial rates. Parentheses represent 95% confidence values from global fits to three independent measurements at each time point. TmA/W: a0 = 2.1 [A-P] (1.9, 2.3), k = 0.5 sec^-1^ (0.3, 0.8), a0*k = 1.1 [A-P]sec^-1^; TmA/W +Tm14FO: a0 = 1.1 [A-P] (0.9,1.4), k = 0.5 sec^-1^ (0.3,1.0), a0*k = 0.6 [A-P]sec^-1^; TmA/W + TarFO: a0 = 0.2 [A-P] (0.1,0.4), k = 0.4 sec^-1^ (0.0,1.3), a0*k = 0.1 [A-P]sec^-1^. (D) Top: Autoradiography of *Tm* CheA autophosphorylation in the presence of CheA, CheW and either the Tm14 or Tar receptor foldon (12 min assay). Bottom: Tar foldon activation of *Ec* CheA (15 sec assay). Samples were run in non-adjacent lanes on the same or parallel-processed gels that were imaged together. See Figure 1 – supplement 4.

### Isolation and analyses of ternary core complexes

MALS was used to find the optimal buffer conditions for producing soluble, homogenous ternary complexes. Complexes composed of either the Tm14 or Tar foldons were isolated by size-exclusion chromatography (SEC) after incubating equimolar amounts of the receptor mimetics with *Tm* CheA and CheW in various buffer conditions. Ternary complex yield increases with buffers of high KCl content (250 mM) and low pH (6.5) (Figure 1 – figure supplement 2). In the case of the Tm14 foldon, subsequent MALS analyses of two resulting high molecular-weight SEC peaks revealed particles of masses 220 kDa and 440 kDa, respectively (Figure 1 – figure supplement 3A,B). Furthermore, these two species differ in their CheA activity, with the larger complex showing more inhibition (Figure 1 – figure supplement 3C). Given the extended nature of the native of arrays, it is unsurprising that the Tar foldon complexes produce several different assembly states; nonetheless, they can be separated based on hydrodynamic properties (Figure 1 – figure supplement 2).

Isolated ternary complexes were analyzed by size-exclusion chromatography coupled to small-angle x-ray scattering (SEC-SAXS) to determine sample size, homogeneity, particle flexibility, and spatial extent. Complexes formed from receptor foldons, CheW, and full-length or CheA domains P3P4P5 represent well-defined particles of expected sizes (Table 1, Figure 1 – figure supplement 3D). SAXS data collected across each respective SEC peak indicates separation of the reconstituted species into relatively homogenous complexes of different sizes (Table 1). In general, the complex size decreases with P3P4P5 compared to full-length CheA, and particle size increases when CheW and the receptor foldons are contained in the complexes.

**Table 1.** Real space SAXS parameters generated from SEC-SAXS for foldon complexes with CheA and CheW.

| Sample | Porod Vol. ($\text{\AA}^3$) | Rg ($\text{\AA}$) | Dmax ($\text{\AA}$) | NSD | Chi<sup>2</sup> | MW (kDa) |
| --- | --- | --- | --- | --- | --- | --- |
| Δ289:CheW (*Tm*) | 266k | 42.6 | 145 | 0.78 | 1.23 | 161 |
| Δ289:CheW:Tm14fo (*Tm*) | 235k | 45.95 | 142 | 0.78 | 1.16 | 190 |
| CheAfl (*Tm*) | 514k | 60.6 | 263 | 0.59 | 1.22 | 250 |
| CheAfl:CheW (*Tm*) | 588k | 57.9 | 238 | 0.74 | 1.16 | 300 |
| CheAfl:CheW:Tm14fo (*Tm*) | 796k | 61.1 | 239 | 0.65 | 1.01 | 355 |
| CheAfl:CheW:Tarfo (*Ec*) | 804k | 58.9 | 226 | 0.58 | 0.95 | 347 |
| Cross-linked<br>CheAfl:CheW:Tarfo (*Tm*) | 796k | 55.4 | 201 | 1.24 | 1.47 | 339 |

Porod Vol.: Porod volume. Rg: Radius of gyration. Dmax: The maximum distance between two points of the particle. NSD: Normalized spatial discrepancy, a quantitative measurement of similarity of three-dimensional envelopes generated from each experiment. MW: Molecular weight of the particle calculated from the Porod Volume.

### Receptor foldons inhibit *Tm* CheA activity

Both the Tar Foldon and Tm14 Foldon were tested for their ability to modulate *Tm* CheA autophosphorylation activity by radioisotope incorporation from *γ*-^32^P-ATP (Figure 1C,D - figure supplement 4). Previous work indicates that Tar receptor foldons activate *Ec* CheA autophosphorylation (Greenswag, AR, et al., 2015); such behavior was recapitulated here and shown to be CheW dependent (Fig. 1D-figure supplement 4A). In contrast, complexes formed by the Tar and Tm14 foldons with *Tm* CheA and CheW deactivate the kinase 20-fold and 2.5-fold, respectively (Figure 1C - figure supplement 4B). At the level of assay detection, the Tar foldon turns off *Tm* CheA activity nearly completely. Given this ability of the Tar foldon to strongly inhibit the *Tm* kinase, Tar foldon complexes with *Tm* CheA and *Tm* CheW (produced at pH 6.5 and 150 mM KCl, Figure 1-figure supplement 2) were isolated for further analysis. To better evaluate the degree to which these particles mimic native assembly states of core signaling particles, we evaluated the consequences of residue substitutions in the PIRs that are known to alter functional properties.

### Receptor foldon residue substitutions affect ternary complex formation

Single residue changes in receptor PIRs affect trimer stability and CheA responses. For example, *Ec* Tsr receptor R388F and R388W (Tar R386F and R386W) and Tsr E391A (Tar E389A) alter cell tumbling bias and trimer stability (Mowery, P, et al., 2008). Tar foldon R386F was significantly more stable than the parent (Figure 1 – supplement 5). This variant can be purified from cell extracts in much higher amounts than WT (20 mg/L of cell culture vs. 6 mg/L), and the purified protein reaches higher concentrations (∼7 mg/ml compared to ∼2 mg/ml; Figure 1 – supplement 5). However, MALS and chemical crosslinking with the lysine-specific chemical crosslinker disuccinimidyl sulfoxide (DSSO) shows that the R386F neither trimerizes nor produces stable complexes with CheA and CheW (Figure 1 – supplement 5). Furthermore, autophosphorylation assays demonstrate that R386F only exhibits ∼1.5-fold deactivation of *Tm* CheA. Similarly, the R386W variant does not trimerize, fails to complex with CheA and CheW, and does not affect CheA autophosphorylation (Figure 1 – supplement 5). Tar foldon E389A is unstable and only present in high-molecular weight aggregates after purification on SEC. Mutagenesis studies and molecular dynamics (MD) simulations implicate the equivalent of Tm14 residue F395 in receptor on-off switching (Ortega, DR, et al., 2013). Tm14 foldon F395W produces similar complexes with *Tm* CheA as the parent, but does not alter kinase activity (Figure 1 – supplement 5). Thus, single residue substitutions in the receptor tips known to affect CheA activity *in vivo* also alter the integrity of the foldon complexes and show perturbed effects on kinase activity, ruling against a largely non-native assembly of the particles.

### Chemical cross-linking of the foldon complexes

We further stabilized the deactivated ternary complex of Tar Foldon, *Tm* CheA, and *Tm* CheW by DSSO cross-linking prior to SEC purification (Figure 2A). The ∼320 kDa MW of the complex (by MALS) agrees with a composition of a core signaling unit (2 Tar foldons, 2 CheA subunits, 2 CheW subunits; Figure 2B). SEC-SAXS of the cross-linked particles produces a MW similar to that found by MALS (MW (V_C_) = 305 kDa; MW (V_P_) = 328 kDa; Table 1, Figure 2 – supplement 1). Some contribution from a higher MW species (∼ 440 kDa) was evident in the SEC-SAXS data when analyzed by singular value decomposition with evolving factor analysis (Meisburger, SP, et al., 2016); however, the larger species component could be separated in the analysis. The dimensionless Kratky plot (Figure 2 – supplement 1) reflects a spherical particle with little indication of the flexibility found in the free CheA kinase (Greenswag, AR, et al., 2015).

**Figure 2.**
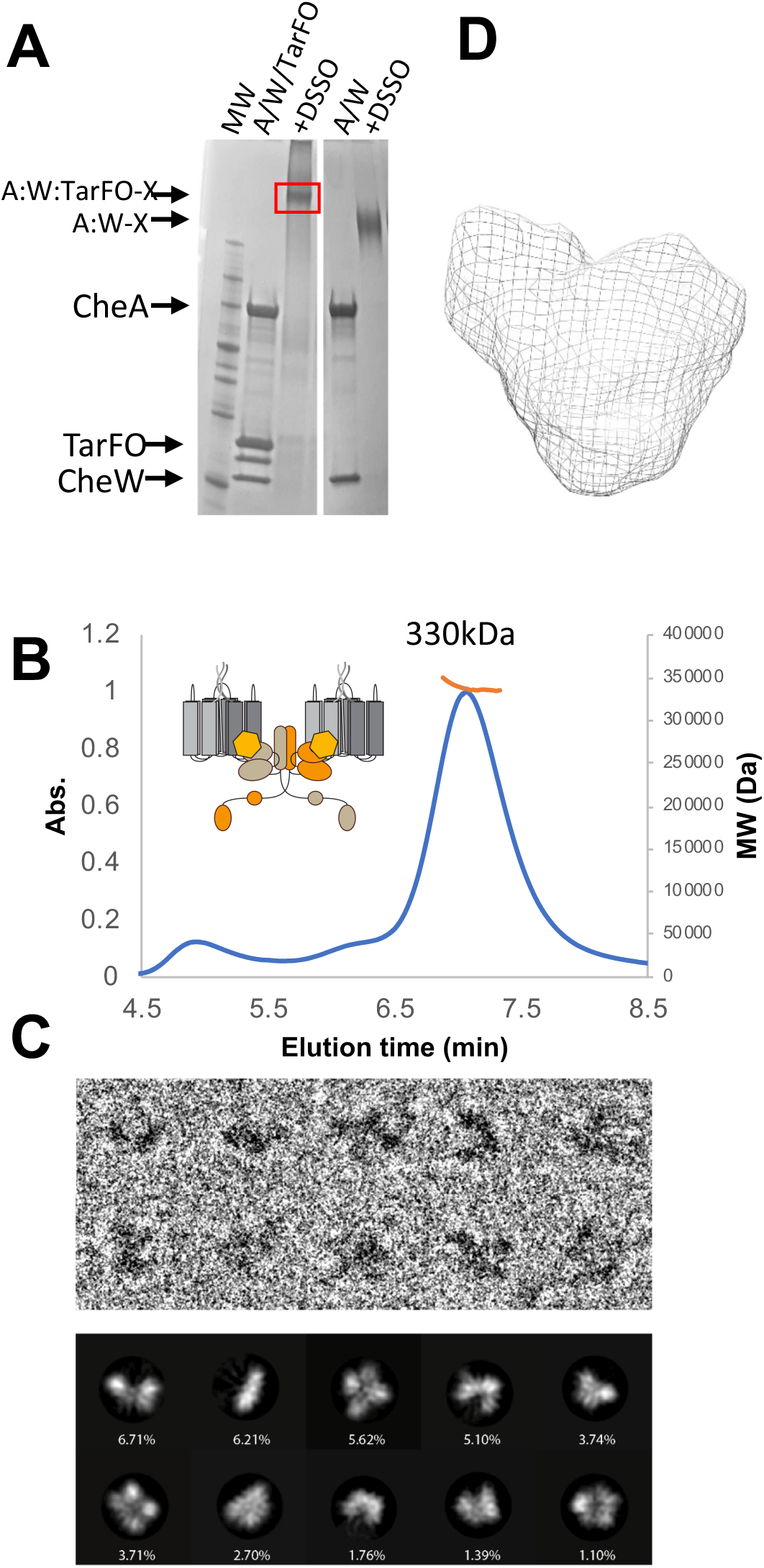
Isolation and analyses of foldon ternary complexes cross-linked with DSSO. (A) SDS-PAGE of the Tar foldon, *Tm* CheA and *Tm* CheW demonstrates the formation of a ternary complex upon addition of DSSO. CheA to CheW DSSO crosslinking is shown on the right. (B) MALS experiments indicate one primary species after SEC purification of the cross-linked complex, MW ∼330 kDa +/- 15 kDa, which would correspond to a core signaling unit containing two receptor foldons, one CheA dimer and two CheW subunits (320 kDa). (C) Cryo-EM of the cross-linked complexes. Upper: Enlarged views of single complex cropped from the micrograph in different orientations corresponding to the class averages shown in lower panel. Lower: Selected 2D class averages and their percentages over the total extracted particles used for alignment and classification (Box size = 268 Å, mask diameter = 140 Å). (D) The molecular envelope of the isolated cross-linked ternary complex generated by SEC-SAXS analyses.

### Cryo-electron microscopy of the foldon complexes

Compositional and conformational homogeneity of the isolated cross-linked complexes was assessed by transmission cryo-electron microscopy (Figure 2C). When unlinked ternary complexes of the Tar foldon with *Tm* CheA and CheW were analyzed, attempts at two-dimensional (2D) classification of the particles were unsuccessful because the particles disassociated on grids and were conformationally heterogeneous (Fig 1 – figure supplement 3E). However, micrographs of the species purified after DSSO cross-linking revealed monodispersed particles and 2D classification generated averages that fit the expected size and shape of the ternary complexes (Figure 2C). Indeed, the 2D classes (Figure 2C) resemble the molecular envelopes constructed from SEC-SAXS experiments in size and general shape (Figure 2D). An insufficient number of suitable particles for analyses currently limits 3D reconstruction of the 2D classes.

### Interface mapping by DSSO-crosslinking and mass-spectrometry

Domain proximity and component interfaces in free *Tm* CheA and in the foldon complexes were analyzed by DSSO crosslinking followed by mass spectrometry. DSSO is a cleavable chemical cross-linker that covalently binds two lysine residues within roughly 20 Å of each other (Kao, AH, et al., 2011). Tandem proteolysis with trypsin and chymotrypsin of the cross-linked samples followed by sequential fragmentation with mass-spec identifies the peptides linked by DSSO. Mass-spec of DSSO-treated *Tm* CheA in the absence and presence of CheW and the Tar foldon reveal key structural changes upon kinase inhibition by the Tar foldon. Among the many cross-links identified (Figure 3 – figure supplements 1,2), the most striking difference between the two samples is that inter-domain cross-links between P1 and P2, P1 and P4, and P2 and P4 are only detected when CheA associates with the deactivating receptor (Figure 3 – figure supplements 1,2). Indeed, the free kinase only produces cross-links *within* P1 or P2 (Figure 3-supplement 1), indicating that these domains are isolated from the rest of the protein. In contrast, the binding of receptor and CheW primarily to P5 associates P1, P2 and P4 to enhance their encounter frequency (Figure 3A and figure supplement 2). The P4 sites that cross-link to P1 and P2 mostly involve residues near the ATP-lid and ATP binding pocket. Furthermore, the cross-linking sites position the His45 substrate residue on P1 away from the P4 ATP-binding site. Instead, the lysine residue nearest His45 preferentially cross-links to the P2 domain rather than the P4 domain (Figure 3A).

**Figure 3.**
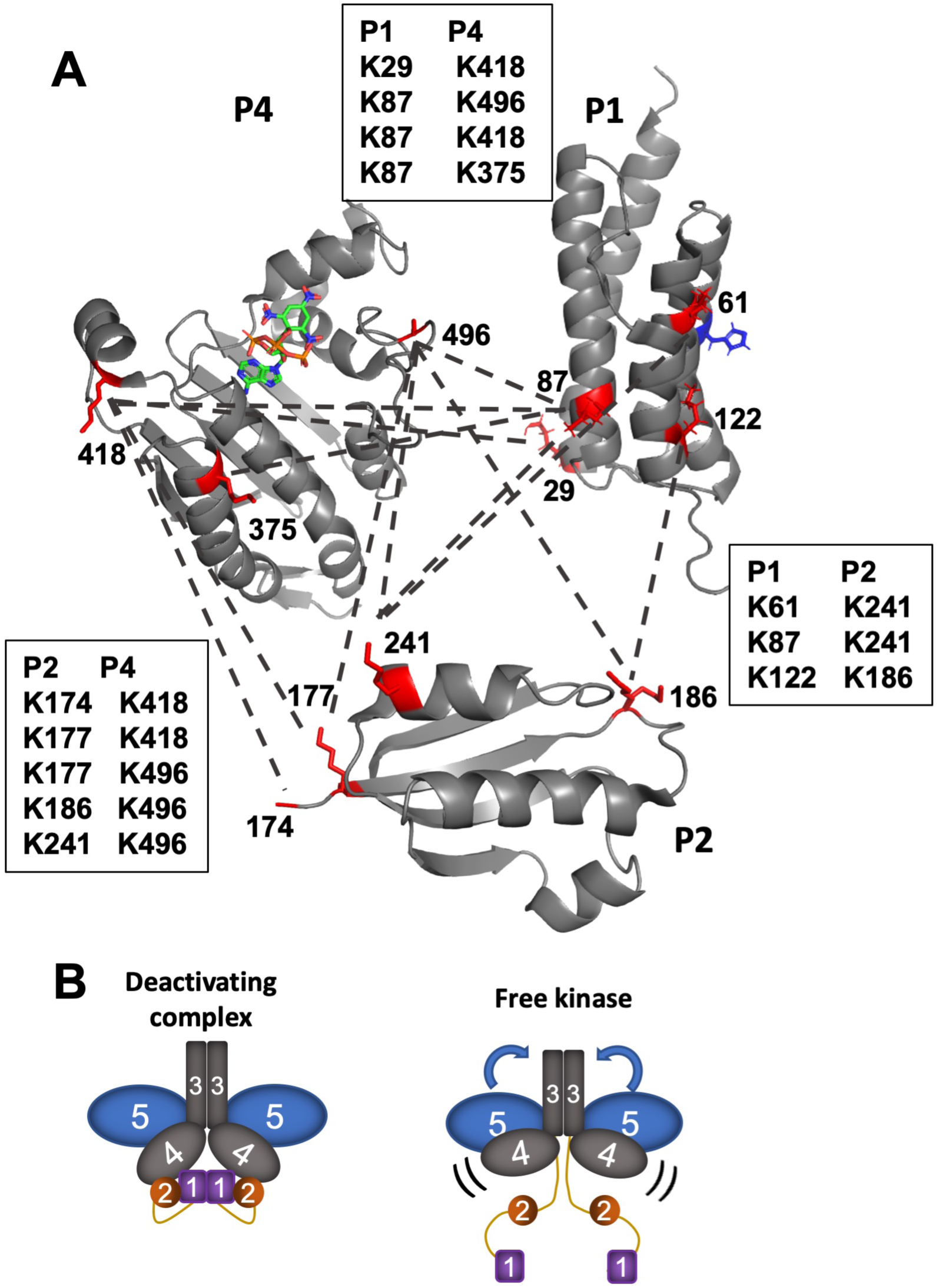
DSSO cross-linking followed by mass spectrometry identifies component interactions in the foldon complexes. (A) Reconstitution of CheA with the deactivating receptor and CheW, promotes interactions among P1 (PDB code: 2LD6), P2 (1U0S) and P4 (1i5D). Cross-linked lysine residues are shown in red and indicated by dashed lines. Residues near helix D on P1, which is distal from the His45 substrate (blue), cross-link to sites near the P4 ATP-binding site. P1 sites near His45 cross-link to residues on the P2 CheY-binding interface. (B) In the free kinase, no inter-domain cross-links were identified for P1 or P2. However, cross-links between P3 and P5 indicate increased flexibility of the P5 domain toward P3 in the absence of the deactivating foldon. In the free kinase compared to the complexes the P4 domains undergo increased inter-domain cross-linking, likely indicative of greater mobility.

Whereas many P1:P2:P4 cross-links are unique to the ternary complex, the CheA-only sample produces contacts between the P3 and P5 domains not found in the foldon complex. Specifically, two lysine residues at the tip of the P3 domain cross-link to P5 lysine residues that are more than 30 Å away in the crystal structure of CheA P345 (PDB code:4XIV). Hence, the P5 domain is highly flexible in the free kinase (Figure 3B). In the ternary complex, the same P3 residues instead cross-link to receptor residues predicted from array structures to be reasonably close (Briegel, A, et al., 2012; Li, X, et al., 2013).

Importantly, inter-protein cross-links identified in the ternary complex are supported by crystal structures of complexes containing CheA P4P5, receptor fragments, and CheW (PDB ID: 3UR1) (Figure 4A-C). CheA P5 and CheW strictly cross-link at binding interface 1 and residues on the upper portion of CheW cross-link to the nearest lysine residue on the Tar foldon (Figure 4A,B). Only lysine residues on the bottom-directed face of CheW cross-link to P4 residues as expected, but these sites are predicted to be more than 30 Å away based on array models (Figure 4C) (Briegel, A, et al., 2012; Cassidy, CK, et al., 2015). Thus, although the P4 domains juxtapose the predicted face of CheW, they retain substantially flexible in the foldon complexes. Nevertheless, taken together, the cross-linking data supports the conclusion that the foldon complexes have structural characteristics expected of core signaling units.

**Figure 4.**
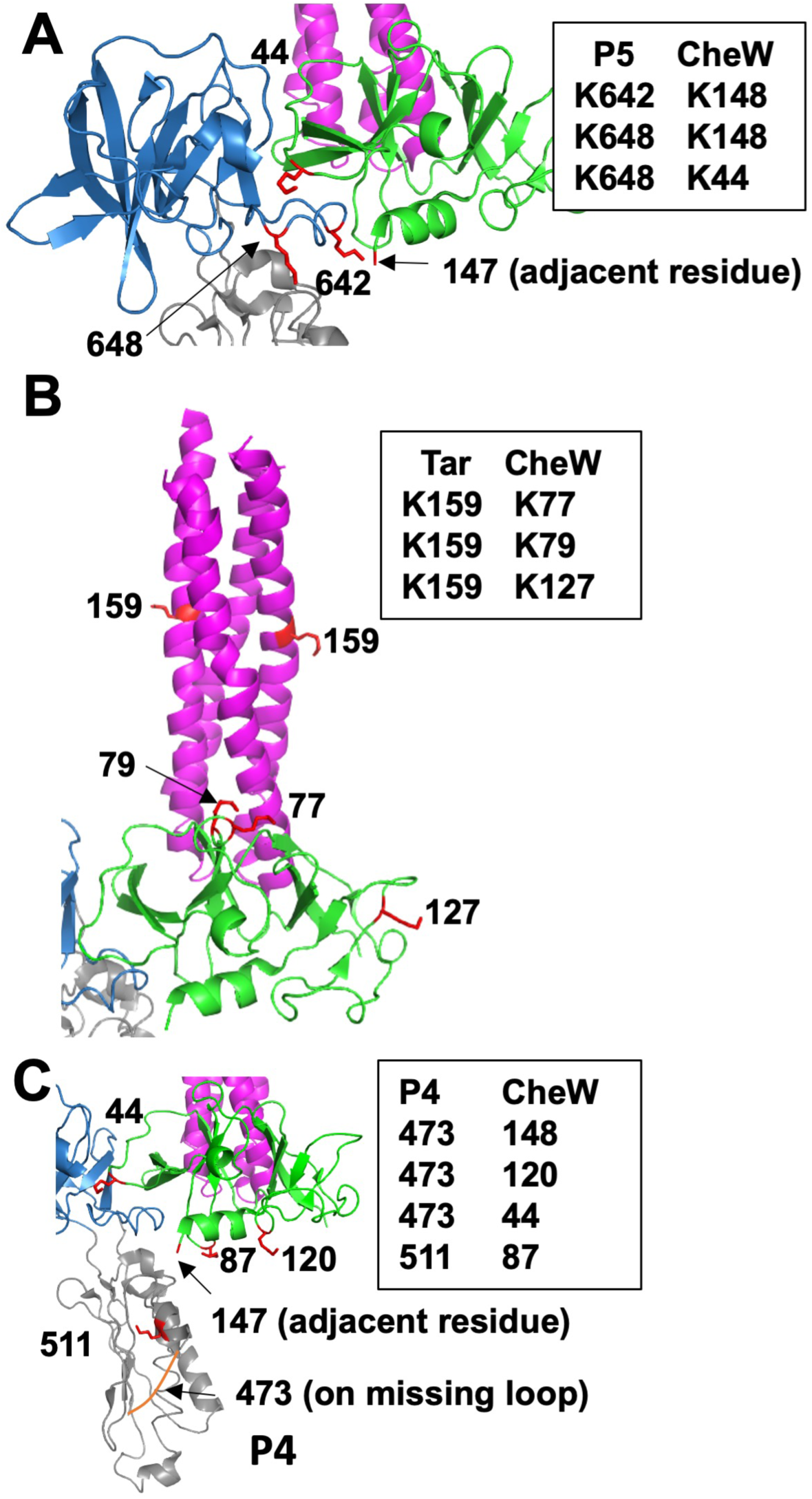
Cross-links identified in the *Tm* CheA, CheW, Tar foldon complex are supported by interaction interfaces in crystal structure 3UR1, which consists of CheA P4P5, CheW and receptor PIRs. (A) The CheA P5 domain (blue) interacts with CheW (green) at lysine residues adjacent to interface 1. (B) Cross-links identified between CheW and the Tar foldon (magenta) indicate that the ‘top’ portion of CheW orients toward the Tar lysine residue that is nearest the receptor tip. (C) Cross-links between the CheA P4 residues near the ATP binding pocket and at the ‘bottom’ portion of CheW indicate that P4 encounters CheW.

### PDS provides evidence for P1 dimerization and P4 dipping within the foldon complexes

Double electron-electron resonance (DEER) spectroscopy of nitroxide-labeled CheA and nitroxide-labeled nucleotide was conducted to characterize CheA domain positioning within the Tar foldon complexes. Nitroxide spin-labels (also known as “R1”) were introduced at residues 12 (P1), 387 (P4), 588 (P5) by cysteine targeting of 1-oxyl-2,2,5,5-tetramethylpyrroline-3-methyl) methanethiosulfonate (MTS) and at the ADP-binding site by use of a previously reported spin-labeled ADP derivative, ADP-NO (Muok, AR, et al., 2018). Each labeling site per subunit generates two R1 moieties in the CheA dimer, between which distance distributions are determined.

In the free kinase, E12C-R1 CheA produced a short (∼28 Å) distance not unlike those observed for previous P1 sites when CheA binds to single-chain “dimeric” receptor fragments (Figure 5A) (Bhatnagar, J, et al., 2010). The short distance becomes somewhat more prominent but does not change dramatically upon incorporation of E12C-CheA into the foldon complex with CheW (Figure 5B). PDS of other P1 sites (E76C-R1, E92C-R1 (Bhatnagar, J, et al., 2010)) also produce distance distributions with maxima in the range of 30-50 Å (Fig. 5B). Interestingly, the crystal structure of the isolated P1 domain (1TQG) from *T. maritima* forms a parallel dimer that is generally compatible with these distance constraints (Figure 6).

**Figure 5.**
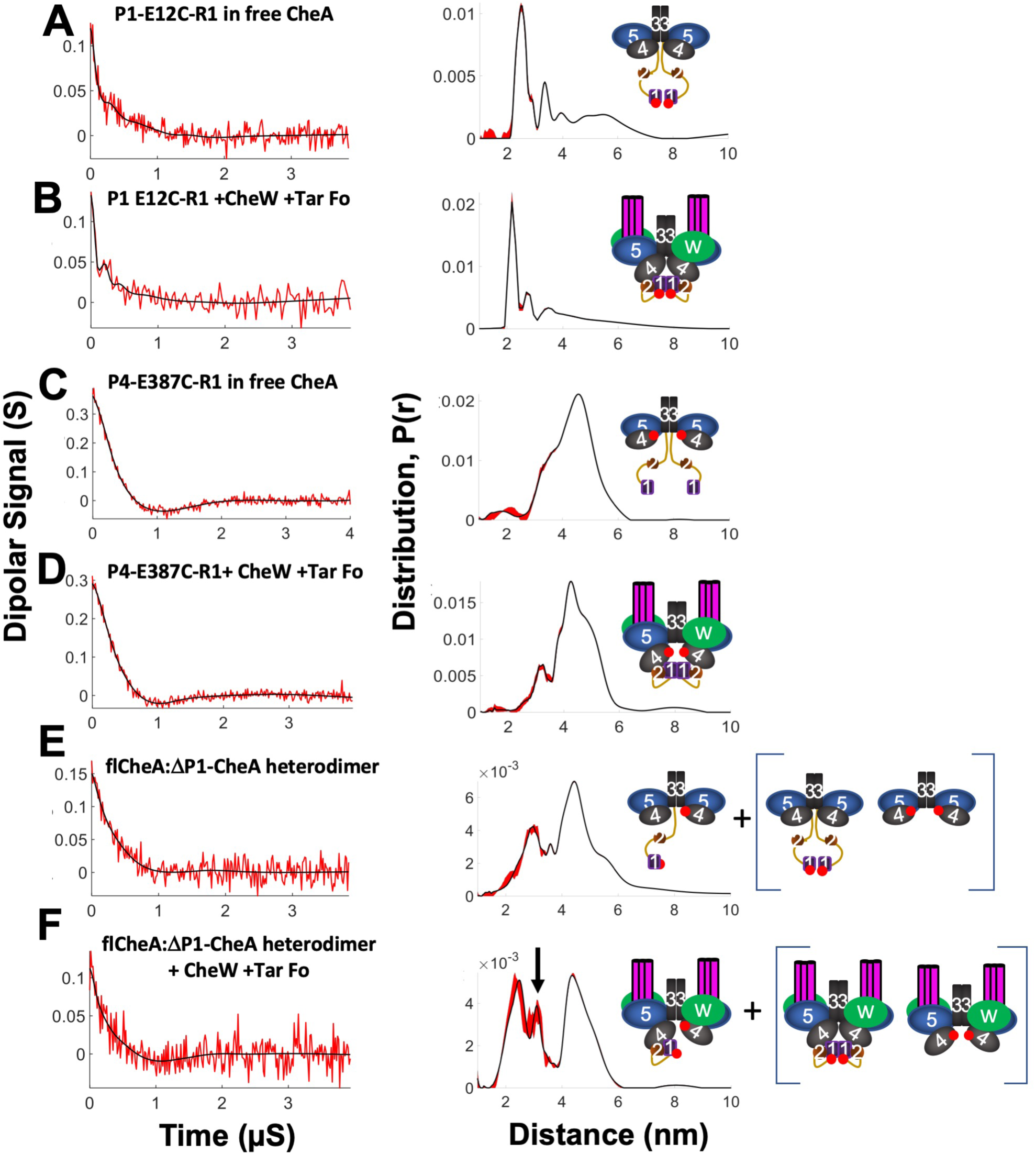
PDS reveals conformational changes in CheA when reconstituted into foldon complexes. All samples were reconstituted with ADP. Schematics at right denote label positions in the context of free CheA or the foldon complexes (red dots represent the R1 nitroxide). Base-line corrected time domain data (left) before (red) and after (black) wavelet denoising and resulting distance distributions (right, black) with error bounds (red) for: (A) P1-E12C-R1 in free CheA; (B) P1 E12C-R1 +CheW +Tar foldon; (C) P4-E387C-R1 in free CheA (D) P4-E387C-R1+ CheW +Tar foldon; (E) flCheA:*Δ*P1P2-CheA heterodimer; (F) flCheA:*Δ*P1P2-CheA heterodimer + CheW +Tar foldon. Arrow denotes new separation arising from presumed P4-to-P1 interaction. Heterodimer samples in (E) and (F) also contain contributions from the homodimeric species shown in brackets.

**Figure 6.**
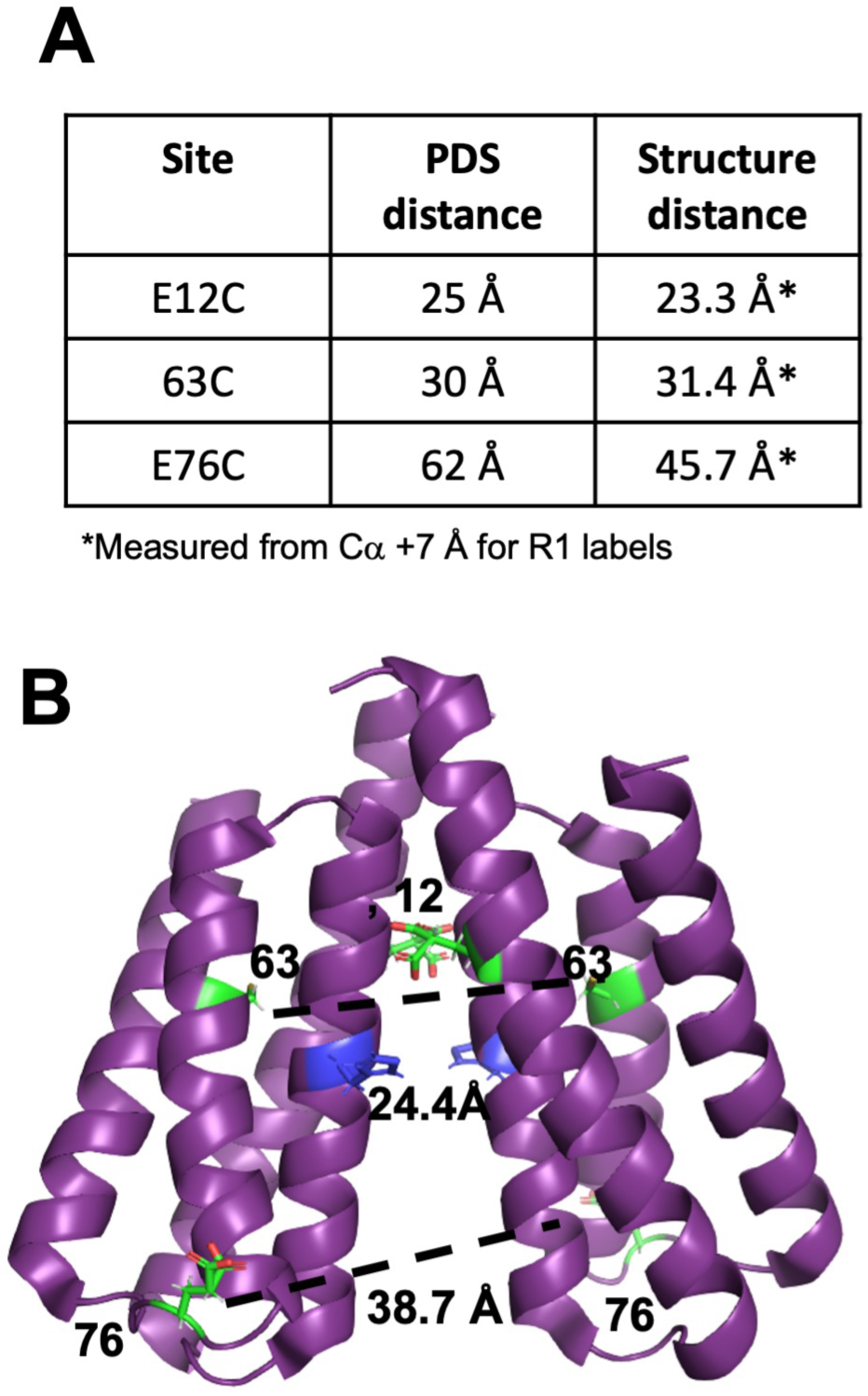
PDS identifies a P1 homodimer in the foldon complex. (A) Summary of distance restraints from DEER experiments on CheA spin-labeled on P1 (this study and Bhatnager et al., 2010). Distances are measured from C*α* positions + 7 Å for R1 labels. (B) The PDS distance distributions generally agree with a parallel P1 crystallographic dimer (1TQG). E76 is located on a flexible loop furthest from the dimer interface. The substrate His45 (blue) resides at the edge of the dimer interface.

The P4 site, CheA E387C-R1, places a spin label at the end of the first P4 *β*-strand, near the L3 linker and projecting toward the CheA dimer interface. As with previous PDS studies of CheA, the broad inter-subunit distance distribution centers at ∼47 Å (Figure 5C) (Bhatnagar, J, et al., 2010; Park, SY, et al., 2006). In the foldon complex, the distance sharpens and a component with a mean separation ∼ 5 Å shorter than that of the free kinase predominates (Figure 5D). To remove contributions from the 387-R1-to-387-R1 interaction and accentuate P1-to-P4 inter-subunit contacts, heterodimers were produced between one subunit of full-length E12C-R1 CheA and one subunit of CheA containing only P3P4P5 E387C-R1 (Park, SY, et al., 2006). Heated incubation of the (flCheA-E12C-R1)_2_ and (P3P4P5-E387C-R1)_2_ dimers encourages subunit swapping and temperature trapping followed by SEC purification enriches for flCheA-E12C-R1:P3P4P5-E387C-R1 heterodimers (Park, SY, et al., 2006). Removal of one P1 domain within the dimer not only dilutes signals from the short P1-P1 interaction but relieves inhibition of CheA autophosphorylation and thus may encourage more P1-P4 contacts (Greenswag, AR, et al., 2015). In the free kinase (Figure 5E) the distance distribution primary shows the ∼47 Å separation typical of the P4-P4 signal from remaining P3P4P5 homodimer and a shorter ∼25 Å separation from full-length CheA dimers also not fully excluded by the SEC purification. There is also some hint of an intermediate distance at ∼36 Å that intensifies in the foldon complex (Figure 5F). This 36 Å signal presumably derives from the interaction between 12-R1 and 387-R1. A 36 Å separation between P1 and P4 agrees well with previous interaction studies (Gloor, SL, et al., 2009; Nishiyama, S, et al., 2014). Models of P1 docked in a productive conformation with the P4 active site (Greenswag, AR, et al., 2015) (Zhang, J, et al., 2005) or a previously characterized inhibitory complex between P1 and P4 (Hamel, DJ, Zhou, H., Starich, M.R., Byrd, A., Dahlquist, F.W., 2006) could produce such a distance.

When treated with a spin-labeled ADP derivative (ADP-NO) that binds to the CheA P4 domain (Muok, AR, et al., 2018) and studied by DEER, the free kinase gives only very weak dipolar signals indicative of long, broad distributions (Figure 7A). In contrast, the foldon complex produces distances at ∼22 Å and ∼35 Å, in addition to some larger (∼50 Å) separations (Figure 7B). The low modulation depth of the time-domain data may derive from the fact that ATP binds *Tm* CheA with substantial negative cooperativity (Eaton, AK, et al., 2009). Thus, even at mM concentrations, the ADP-NO is likely not bound in both subunits for a substantial fraction of the sample. Nonetheless, the short distances indicate that the Tar foldon causes the P4 domains to move together from their positions in free CheA. The bimodal distribution indicates that either the P4 domains or the nitroxide moieties themselves exhibit several related conformations. To further bound the ATP-binding site position, CheA was labeled on the P5 domain at position 588 of subdomain 2. Free CheA E588C-R1 gives nearly no dipolar signals in keeping with the predicted long (> 80 Å) separation between these sites. However, in the foldon complex, a very short distance component arises from E588C-R1 (Figure 7 – supplement 1). Similar short distance components on the spin-labeled P5 domain are known to derive from CheA dimer association through symmetric subdomain 2 contacts (Bhatnagar, J, et al., 2012). Such contacts mimic those of interface 2 in crystal structures and arrays, but in this case, are likely favored by the high-glycerol and low temperature conditions of the DEER experiments. Interestingly, binding the Tar foldon and CheW further increases these inter-dimer contacts relative to the free kinase (Figure 7 – supplement 1), probably because the receptors align the P5 domains in planes compatible with symmetric binding at the P5 domain ends. Addition of ADP-NO to free CheA E588C-R1 produces some shorter distances at ∼45 Å, but mainly large distances (> 60 Å) that represent the P5- to-P4 separation are evident (Figure 7C). In contrast, the Tar foldon complex with E588C-R1 and ADP-NO (Figure 7D) produces a mid-range distance at ∼50 Å and a short distance at ∼20-25 Å that matches overlapping contributions from the inter-dimer contact (Figure 7 – supplement 1) and the ADP-NO contributions seen in absence of E588C-R1 (Figure 7B). The 50 Å distance, which intensifies relative to the long-distance component seen with the free kinase (Figure 7C), likely reflects an ordering and contraction between P5 subdomain 2 and the ATP binding pocket of P4 (Figure 7D). Longer distances (> 60 Å) may represent greater separations of these units, as observed in the free kinase. In summary, the Tar foldon complexes enforce a kinase conformation largely consistent with array models. Importantly, the PDS data also provides direct evidence for the close interaction between the P4 domains that has been predicted by MD simulations (Cassidy, CK, et al., 2015; Cassidy, CK, et al., 2020).

**Figure 7.**
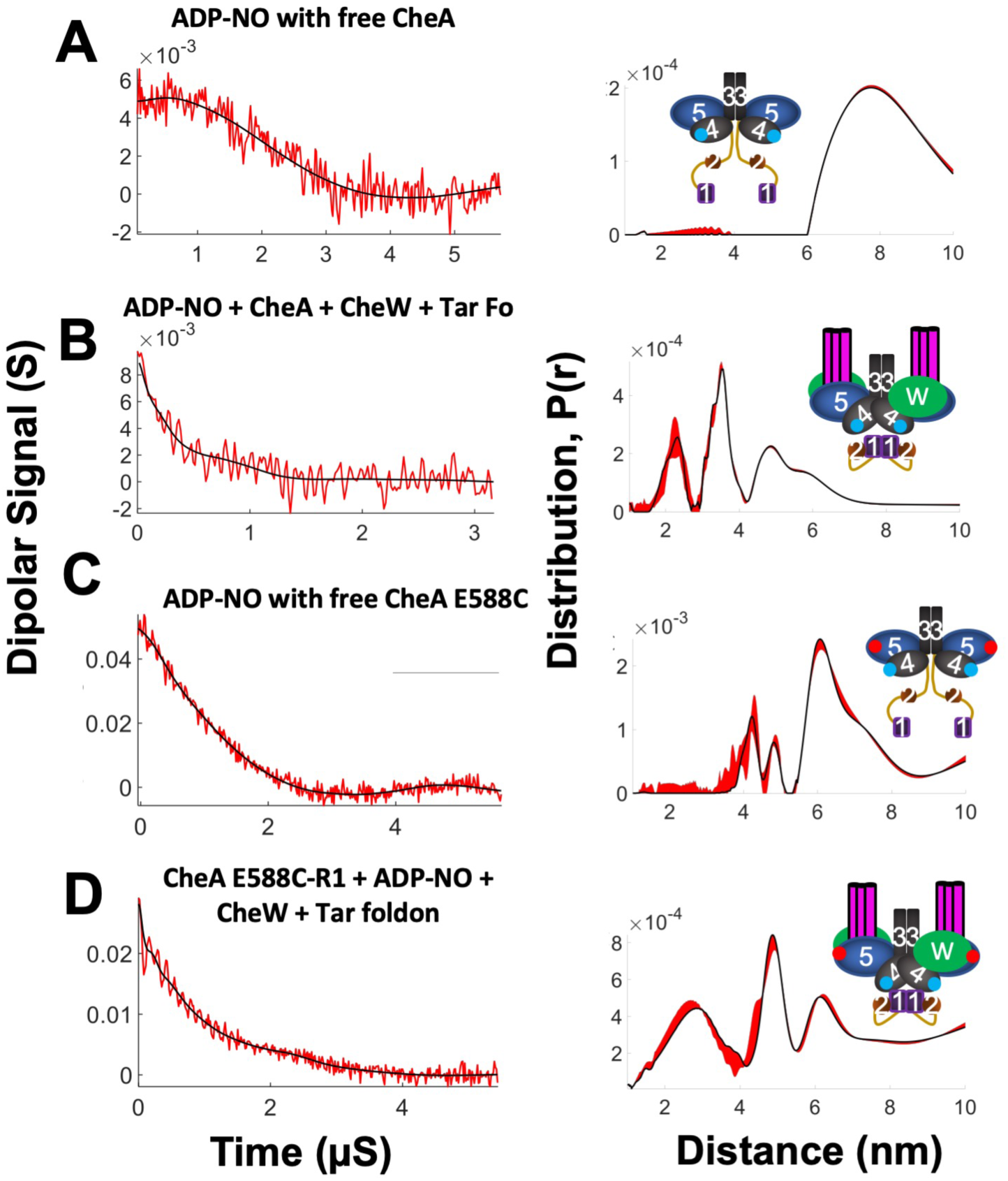
PDS with spin-labeled ADP-NO track the CheA ATP-binding pocket. Schematics at right denote label positions in the context of free CheA or the foldon complexes (red dots represent R1 nitroxide, blue dots represent ADP-NO). Base-line corrected time domain data (left) before (red) and after (black) wavelet denoising and resulting distance distributions (right, black) with error bounds (red) for: (A) ADP-NO with free CheA.; (B) ADP-NO, Tm CheA, CheW and Tar foldon; (C) ADP-NO with free CheA E588C; (D) CheA E588C-R1 with ADP-NO, CheW and Tar foldon.

### Modeling of the foldon complexes

Stepwise modeling of the foldon core complexes was based on their determined component stoichiometries, SAXS data, cross-linking constraints, PDS restraints, and molecular structures determined from crystals and arrays in native and reconstituted forms (Figure 8). Cryo-ET indicates that in arrays the CheA P4 domains assume a different conformation relative to the P3 domains than found in the structure of CheA P3P4P5 (Bilwes, AM, et al., 1999), but may conform more closely to the positions observed in the crystal structure of CheA P3P4 (Greenswag, AR, et al., 2015). Thus, P4 was suspended below the P5-CheW layer based on the P3P4 structure (4XIV) and the L3 and L4 linkers were removed and rebuilt using the Rosetta KIC (Kinematic Closer) loop modeling algorithm. Allowable backbone torsion angles, bond angles and bond length were calculated, and L3 and L4 linker connections were built consistent with their spatial and sequence constraints. The highest scoring CheA model was associated with CheW and receptor trimers based on known crystallographic interfaces and was then minimized against stereochemical and packing functions of the Rosetta Relax program with additional distance restraints from the PDS data. The output models were then ranked based on their Rosetta all-atom scores. The P1 domains were dimerized based on the crystallographic structure and PDS restraints and situated below the P4 domains to align the molecular symmetry axes. This configuration is generally consistent with cryo-ET density tentatively assigned to the P1 and P2 domains (Briegel, A, et al., 2013), as well as the spatial extent of the SAXS envelopes generated for the cross-linked complex (Figure 8). The P1 dimer position matches sites on P1 and P4 involved in a previously characterized inhibitory contact (Hamel, DJ, Zhou, H., Starich, M.R., Byrd, A., Dahlquist, F.W., 2006). The P2 domains where then situated beside the P4 and P1 domains to better fill the SAXS envelopes and align the molecular faces that undergo interdomain cross-linking. Domain orientations were chosen to minimize distances given by DSSO cross-linking (20 ± 7 Å). The ADP-NO inter-subunit and P5-to-ADP-NO distances provide strong evidence for dipped P4 and the Rosetta modeling confirms that such conformations can be obtained with reasonable L3 and L4 geometries. Although placement of the P1 domains and especially the P2 domains are not precisely constrained by the cross-linking data, these domains clearly co-localize with P4 in the inhibited complex and Lys residues on specific faces show preferred cross-linking patterns (Figure 8). To produce better agreement of the model with the SAXS data for the cross-linked particle, the L1 and L2 loop conformations were first built automatically as C*α* traces, elaborated as full polypeptide chains and then evaluated for consistency with the SAXS scattering curves. Configurations that placed the large L1 linker below the P1 domains and the L2 linkers to the side of P4 produced the best agreement; conformations related to these were not significantly distinguished by the SAXS data. The entire complex, including CheW and the foldon receptors was then relaxed in Rosetta under constraints of the ESR data and the cross-linking distances. One hundred new models were generated and each was evaluated for agreement against the SAXS data (Figure 8 – supplement file 1A). In the final step, the best model was then further optimized against the SAXS data by configurational sampling about the P4-P5 linkage (P5-CheW and receptor foldons were treated as rigid bodies) to allow for flexibility in the absence of the constraints provided by native membrane insertion (Figure 8 – supplement file 1B). Neither multi-state modeling nor treating the L1 and L2 loops as highly disordered segments significantly improved fits to the SAXS data. The current working model (Figure 8) agrees reasonably well with the SAXS scattering data (*χ*^2^ = 4.5; Figure 8 – supplement 1C), matches the size and shape of the SAXS-derived molecular envelope and is consistent with current distance restraint data (Figure 8 – supplement 2). Although the model is not a unique high-resolution solution to the current data, it encapsulates definitive structural features of the particle including: interfaces among P5, CheW and the PIRs defined by previous crystal structures, movement of the P4 domains toward one another relative to the free kinase, association of the P1, P2 and P4 domains, dimerization of P1 between the P4 domains and localization of the L1 linker below the P1,P2 and P4 cluster.

**Figure 8.**
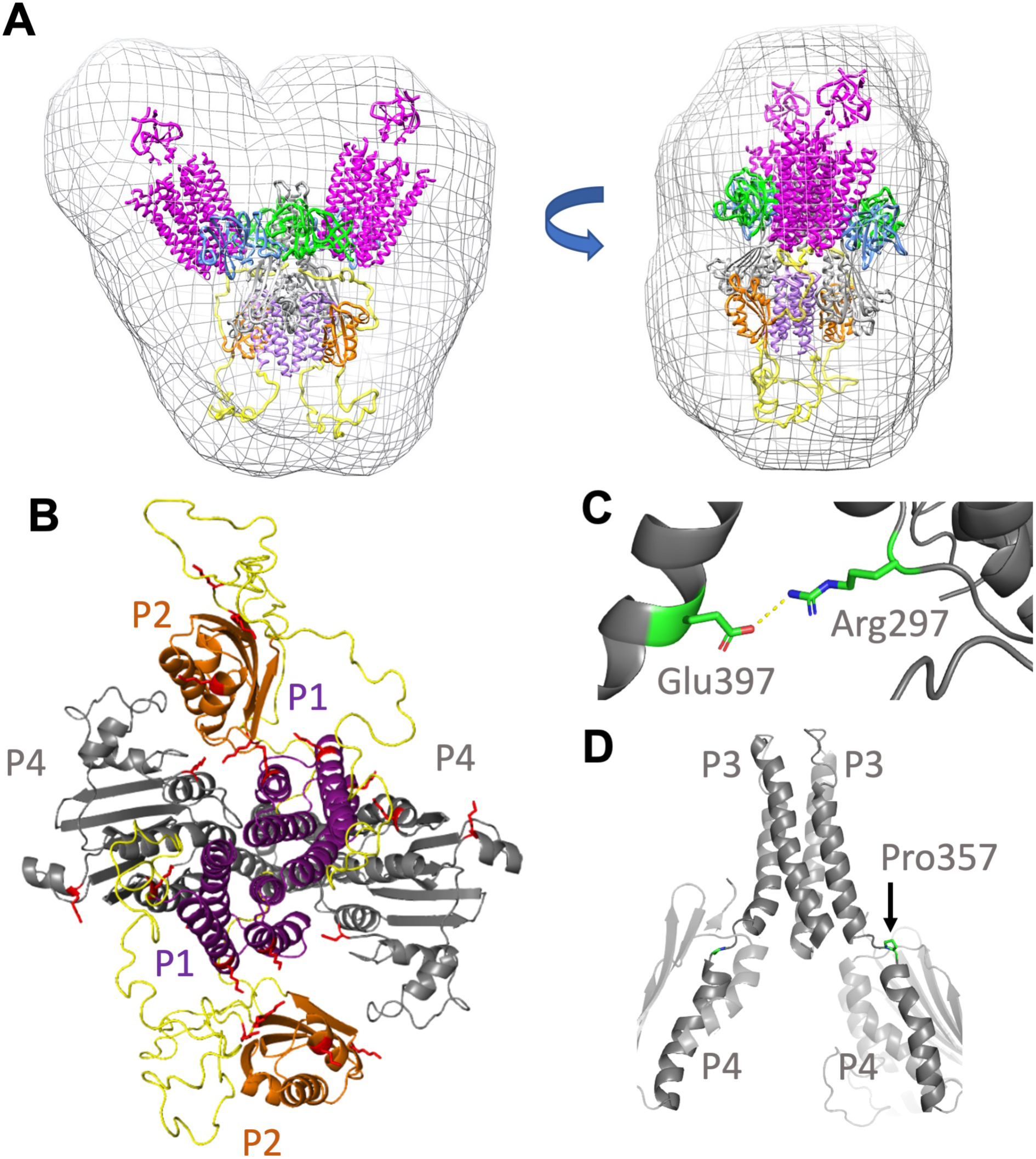
Model of the ternary complex derived from crystal structures, PDS, SEC-SAXS and cross-linking. (A) The working model fits the size and shape of the molecular envelope generated by SEC-SAXS of the cross-linked ternary complex. P1: purple, P2:orange, P3-P4:grey, P5:blue, CheW:green, L1-L2:yellow (B) The lysine residues identified in DSSO cross-links among the P1, P2 and P4 domains (red) localize to internal interfaces in the ternary complex model. (C) Salt bridge between residues Glu397 and Arg297 (shown in Chain A) has been implicated with receptor-coupled CheA activity and are maintained in the working model. (D) The L3 linker between the CheA P3 and P4 domains in both subunits forms a discontinuous helix (Pro357 sidechains highlighted) in the highest scoring Rosetta model.

## Discussion

The receptor foldons interact with CheA and CheW to form complexes that can be isolated in milligram quantities as homogenous ternary complexes. Stability and uniformity of the complexes improves through chemical cross-linking. The resulting cross-linked particles mimic a core signaling unit (2 TODs, 2 CheA subunits, and 2 CheW subunits), which is the minimum component of the chemosensory array that maintains modulation of CheA kinase activity and some cooperative responses (Li, M, et al., 2014; Li, MS, et al., 2011).

Thus, these particles are suitable for examining the domain arrangements of CheA in a receptor-regulated state; in particular, complexes of *Tm* CheA with foldons derived from the *Ec* Tar receptor produce an especially stable kinase-off configuration. Expressed in cells or reconstituted *in vitro* with CheA and CheW, MCP cytoplasmic fragments (C-frags) are well known to affect CheA activity (Airola, MV, et al., 2013; Ames, P, et al., 1994; Asinas, AE, et al., 2006; Levit, MN, et al., 2002; Montefusco, DJ, et al., 2007; Shrout, AL, et al., 2003; Surette, MG, et al., 1996). Some in vitro complexes highly activate CheA (Levit, MN, et al., 2002; Surette, MG, et al., 1996) and produce relatively homogeneous particles that are larger than a core unit (Wolanin, PM, et al., 2006). C-frag complexes of CheA and CheW characterized by pulsed ESR spectroscopy (Bhatnagar, J, et al., 2010), hydrogen-deuterium exchange mass-spectrometry (HDX) (Koshy, SS, et al., 2014) and solid-state NMR (Harris, MJ, et al., 2016) reveal aspects of component interactions but have not provided holistic views of the core signaling unit. His-tagged mediated dimerization (Haglin, ER, et al., 2017) and templating of C-frags on lipids (Cassidy, CK, et al., 2015; Shrout, AL, et al., 2003) have been useful strategies for analysis by cryo-ET (Cassidy, CK, et al., 2015). The foldon approach leverages trimerization of the single chain C-frag fusions to produce particles of well-defined size that contain components of a single core module.

Cross-linking mass-spectrometry experiments on the inhibited CheA foldon complex support a model where the CheA P1, P2 and P4 domains localize in the deactivated kinase. This result is consistent with cryo-ET studies of receptor variants that deactivate CheA and show enhanced “keel” density below the P4 domains that depend on the presence of P1 (Briegel, A, et al., 2013; Yang, W, et al., 2019). Importantly, inter-domain cross-links between P5 and CheW, as well as between CheW and the receptor, match expectations from crystal structures and array models, thereby indicating that the foldon complexes recapitulate functionally relevant contacts among the components. Although cross-linking reveals interactions among P1, P2 and P4 in the deactivated state, no such interactions are found without receptor foldons and CheW. Association of *Tm* CheA with TODs indirectly co-localizes these domains because the receptors primarily interact with CheW and P5, and do not likely contact P1, P2 or P4. The inter-domain contacts between P1 and P4 orient P1 helix D toward the ATP-binding site of P4. Anchoring P1 here may contribute to kinase inhibition by shielding His45 from P4. Indeed, previous NMR-shift assays of P1 binding to P3P4 identified interaction sites between helix D of P1 and a cluster of residues close to the L3 linker on the C-termini of both *α*1 and *β*1 of P4 (Hamel, DJ, Zhou, H., Starich, M.R., Byrd, A., Dahlquist, F.W., 2006). The cross-linking pattern of the inhibited foldon complex is consistent with this interaction mode and the resulting model places the P4 *β*1 site directly in the interface with P1 helix D and the *α*1 site in contact with the L3 linker that changes conformation upon P4 dipping. The PDS measurements surprisingly reveal P1 dimerization, as found in the *Tm* P1 crystal structure (Figure 6). The P1 dimer occludes access of the P4 ATP pocket to His45, but leaves helix D available for interaction with P4. Whereas interactions between P1 and P4 in the inhibitory complex were reasonably anticipated, interactions among P1, P2 and P4 were not. For phosphotransfer to CheY, P2 need not interact with P1 or P4, yet P2 preferentially cross-links to these domains and not to P5, CheW or the receptor foldons. Interestingly, P2 sites on the CheY interaction surface (Park, SY, et al., 2004) cross-link to a Lys residue near the P1 substrate His45. Thus, P1-to-P2 contacts could potentially interfere with both His45 phosphorylation and CheY binding. P1-P1 interactions are also supported by the observations that the P1 domains cross-link to each other (Greenswag, AR, et al., 2015) and increase the subunit affinity of the CheA dimer (Kott, L, et al., 2004). P1 dimerization that reduces His45 accessibility may also explain why both *Ec* and *Tm* CheA activity increases upon removal of one P1 domain (Greenswag, AR, et al., 2015; Levit, M, et al., 1996).

Compared to their mobility in the free kinase, the P4 domains order and move toward each other in the foldon complexes. MD simulations of the core signaling particle composed of CheA P3P4P5 capture transitions between a CheA conformation in which the P4 domain interacts with the P5-CheW layer (undipped) and a conformation where P4 moves down toward the adjacent P4 domain (dipped) (Cassidy, CK, et al., 2015; Cassidy, CK, et al., 2020). Positioning of the P4 domains based on the ADP-NO interaction and the E387C-R1 contact combined with Rosetta loop modeling of the L3 and L4 linkers provide strong support for the dipped conformation of P4 in the inhibitory foldon complex. These configurations are consistent with recent high resolution cryo-ET density of the P4 region in reconstituted arrays on lipid supports that contain CheA P3P4P5; however, disorder in the cryo-ET density makes precise P4 positioning difficult (Cassidy, CK, et al., 2020). The MD simulations further suggest that key salt bridges between the P3 domain (*Ec* Arg265) and P4 residues (*Ec* E368,D372) stabilize the dipped conformation (Cassidy, CK, et al., 2015; Cassidy, CK, et al., 2020). Residue substitutions at these positions affect CheA activity and chemotaxis (Pinas, GE, et al., 2019). In particular, charge reversals at several key sites substantially reduce receptor-coupled CheA activity and regulation by attractant (Pinas, GE, et al., 2019). Thus, it has been suggested that the active state of CheA involves a dipped conformation (Pinas, GE, et al., 2019). The MD simulations also indicate that formation of a nearly continuous helix between P3 *α*2 and P4 *α*1 stabilizes the dipped conformation. The Rosetta data-restrained models also recapitulate the salt bridge between *Ec* Arg265 (*Tm* R297) and *Ec* E368 (*Tm* E397; Figure 8C); however continuous helicity breaks at *Ec* Val344-Pro345 (*Tm* V356-P357) and the P3 and P4 helices are not fully aligned (Figure 8D).

Despite harboring a “dipped” P4 conformation, CheA autophosphorylation activity is low in the foldon complexes, at least in part because both P1 and P4 are sequestered. Could then a dipped conformation be associated with both an inhibited and activated state of CheA? Several related conformations that all suspend P4 away from the P5-CheW layer could be collectively important for both the inhibited and activated states. Both P4-P4 cross-linking, which decreases in the foldon complexes compared to free CheA, and the PDS distances, rule against direct P4-P4 contacts in this particular inhibited state. However, negative allostery in nucleotide binding and the low activity of P4 monomers for P1 phosphorylation (Greenswag, AR, et al., 2015; Hamel, DJ, Zhou, H., Starich, M.R., Byrd, A., Dahlquist, F.W., 2006) suggest that the P4 domains do influence each other either directly or indirectly. When the P1 domains release to become substrates, the P4 domains may further approach one another. As has been suggested (Pinas, GE, et al., 2019), transitions of P4 among conformational states may be key for the catalytic cycle. Under this assumption, restricting such motion, through interaction with P1 or possibly the P5-CheW layer, would provide the means to curtail autophosphorylation. Whether the key regulatory conformations also involve a fully undipped conformation is unclear. Undipped P4 may allow for interaction of the ATP-binding regions with the P5-CheW layer, but it may also be favored by the absence of the P1 and P2 domains in the reconstituted arrays and simulations.

Thus, in the *Tm* inhibited foldon complex, the P4 domains are likely juxtaposed and the P1, P2 and P4 domains are closely associated in a region distal from the P3, P5, CheW, and receptor components. Receptor binding to remote sites on P5 and CheW promote these interactions. In consideration of these findings it is important to note that core complexes composed of CheA P3P4P5 display normal receptor-mediated inhibition of CheA activity, even with free P1 as the substrate (Mello, BA, et al., 2018; Pan, W, et al., 2017). Thus, neither covalent attachment of P1 nor the presence of P2 is required for receptor-regulated kinase inhibition. Furthermore, saturating levels of free P1 do not curtail kinase activity (Mello, BA, et al., 2018; Pan, W, et al., 2017). Nonetheless, a P1 inhibitory site could be operative in the kinase-off state, if kinase activation then renders this site unavailable. Changes in free P1 phosphorylation manifest from changes in the catalytic rate constant (k_cat_) not the Michaelis constant (K_M_) (Mello, BA, et al., 2018; Pan, W, et al., 2017). Thus, kinase inhibition may derive from a depopulation of CheA states capable of productive interactions with P1. A bound P1 dimer may contribute to stabilization of an inhibited state, provided such interactions do not depend on covalent attachment to the core complex and they are relieved in the kinase-on state. The structure of the foldon receptor complexes prevents access of the P4 ATP binding pocket to the P1 substrate His. Reduced domain dynamics in the foldon complexes, communicated from the receptors to P4 through its linkers to P3, or through interactions with the P5-CheW layer, may favor P1 binding in this non-productive configuration.

Indeed, increased order in the arrays, especially in those CheA regions suspended below the P5-CheW layer appears to be a hallmark of the inactive state (Yang, W, et al., 2019). Initial studies of CheA activity in membranes descripted a receptor-inhibited state that was designated as “sequestered” because it could not autophosphorylate P1 nor exchange phosphate with ADP once phosphorylated (Borkovich, KA, et al., 1990). The data presented here indicate that an inhibited core signaling unit sequesters both the P1 and P4 domains through their non-productive interaction and that alleviation of these restraints would be required for kinase activation.

## Methods and Materials

### Cloning and protein purification

All proteins were cloned into pet28a vector containing a kanamycin resistance marker and transformed into *E. coli* (DE3) competent cells. After plating the transformation on agar containing kanamycin, a single colony was chosen for cell culture growth. 8 liters of cells were grown at 37°C until an O.D. at 600 nm reached 0.6. 1mM IPTG was added to each flask and the cells were grown for 16 hours at room temperature. After pelleting via centrifugation, the cells were resuspended in 50 mL of lysis buffer (50 mM Tris pH 7.5, 150 mM NaCl, 5mM Imidazole) and sonicated for 6 minutes. The lysed cells were then centrifuged at high speed to remove the insoluble fraction from the lysate. The lysate was then run over Nickel-NTA affinity resin, washed with 20 mM imidazole buffer to remove non-specific binding proteins, and finally eluted using buffer with 200 mM imidazole. The eluted protein was further purified by size-exclusion chromatography (prep-grade Sephadex S200 or s75). The fractions containing the protein of interested were pooled and concentrated via high-speed centrifugation. The purification protocol was conducted at 22 °C for CheA and CheW and 4°C for the receptor foldons.

### Phosphorylation assays

25 µl samples containing 2 µM CheA (subunit) were incubated in the presence or absence of 2 µM CheW and 6 µM receptor foldon (subunit) in 50 mM MOPS pH 7.5, 150 mM KCl, 10 mM MgCl_2_. Phosphorylation of CheA was initiated by the addition of 1 mM ATP mixed with radiolabeled γ-^32^P ATP at a final radioactivity of 0.15 ± 0.05 mCi and quenched at 1-12 min for *Tm* CheA and 15 s for *Ec* CheA with 25 µl of 3X LDS buffer containing 100 mM EDTA. The samples were run on a native Tris-glycine gel for 2 hours at 120 volts. The gels were dried, placed in a radiocassette for 24 hours, and then imaged with a Typhoon phosphor-imager. Band intensities were quantified using Image J software. Autophosphorylation kinetics were fit to a first-order model of the form a = a_0_(1-exp^-kt^), where k is the first order rate constant and a_o_ represents the final relative level of autophosphorylation.

### Multi-angle light scattering

Reverse-phase chromatography coupled to multi-angle light scattering experiments was used to determine the molecular weight of isolated ternary complexes. For each sample, 2-5mg/ml of protein was injected onto a Phenomenex reverse-phase column pre-equilibrated with 50mM Tris pH 7.5, 150 mM NaCl at room temperature. Bovine serum albumin (sigma) was used as a protein standard. Wyatt technologies ASTRA 6 program was used for data analysis and molecular weight calculations.

### Preparation of isolated ternary complexes

Ternary complexes consisting of receptor foldon, CheA and CheW were produced in milligram quantities by mixing the composite purified proteins in a 1:1:1 stoichiometric subunit ratio in 50 mM Tris or HEPES pH 7.5, 250 mM KCl, 5mM MgCl_2_, 10% glycerol and allowing the mixture to incubate at 4°C for at least 1 hour. After incubation, the sample is injected onto two prep-grade Sephadex columns (s200 and s300) attached to run in tandem and the elutant was collected in 6 mL fractions. Protein components in each fraction were evaluated by SDS-PAGE. Fractions of interest were concentrated to ∼ 8 mg/ml via high-speed centrifugation, flash frozen, and stored at −80°C.

### Size-exclusion chromatography small-angle x-ray scattering (SEC-SAXS)

SAXS data were collected at CHESS G1 line on the Finger Lakes CCD detector. For each sample, ∼2-3 mg/ml of protein was injected onto a Superdex 5/150 size-exclusion increase column pre-equilibrated with sample buffer (50 mM Tris pH 7.5, 250 mM KCl, 1 mM MgCl_2_, 4% glycerol) coupled to the G1 SAXS sample cell. The eluted sample was collected with 2 second exposure times to the x-ray beam at a flow rate of 0.15 ml/min. AMBIMETER (Svergun, DI, 1992) and GNOM (Semenyuk, AV, et al., 1991) were used to initially assess the quality of the scattering data and particle homogeneity. RAW (Hopkins, JB, et al., 2017) and PRIMUS (Konarev, PV, et al., 2003) were used for processing of the SEC-SAXS data and to generate Kratky plots. Further analysis including dimensionless Kratky were carried out with ScÅtter (Rambo, RP, et al., 2013). Background scattering was determined from SEC fractions containing primarily buffer. Molecular weight estimates were made from the Porod volume (V_P_), volume of correlation (V_C_) (Rambo, RP, et al., 2013) and from envelope reconstructions. Envelope reconstructions were calculated with DAMMIF (Franke, D, et al., 2009) and DAMAVER (Volkov, VV, et al., 2003). Ten models were independently generated and then averaged into a consensus envelope without assuming particle symmetry. For the cross-linked ternary complex singular value decomposition with evolving factor analysis (Meisburger, SP, et al., 2016) was applied to separate the minor contribution of a higher MW species from the leading edge of the SEC curve. Model agreement to SAXS scattering curves was evaluated with FoXS (Schneidman-Duhovny, D, et al., 2016).

### Cryo-electron microscopy and image analysis

For cryo-specimens, 3 µL aliquots of 0.1 mg/ml cross-linked ternary complexes were applied to freshly glow-discharged R2/2 Quantifoil 200 mesh copper grids (Quantifoil Micro Tools). Grids were blotted in a climate chamber set to 20 °C and at 95% humidity before plunge frozen in liquid ethane set at −183 °C using a Leica EM GP system (Leica Microsystems). Images containing complexes were collected manually with a 120 kV TALOS transmission electron microscope (Thermo Fisher Scientific) at a magnification corresponding to a pixel size of 1.4 Å in a defocus range between 2 to 4 microns. Image analysis was done with RELION −3.0.2 using ∼ 34,000 particles from 65 images. 2D classification results were generated with a selection of ∼ 22,000 particles.

### Protein cross-linking

Crosslinking of ternary complexes was accomplished by first incubating the receptor foldons with CheA and CheW in a 1:1:1 stoichiometric ratio to a final concentration of 10 µM of each protein in 50 mM HEPES pH 7.5, 250 mM KCl, 5 mM MgCl2, 100 µM ADP, 10% glycerol. After incubation for at least 30 minutes at 4°C, the chemical crosslinker DSSO is added to the protein mixture to a final concentration of 1 mM and crosslinking proceeds for 30 min to 1 hour at room temperature. The cross-linking reaction was quenched by the addition of Tris pH 8.0 to a final concentration of 20 mM. The reaction mixture is run through size-exclusion chromatography on two prep-grade sephadex columns (s200 and s300) attached to run in tandem with a 6 ml fraction volume. Fractions of interest were selected to for further analysis by SDS-PAGE and MALS.

### Mass spectrometry of cross-linked protein complexes

#### Sample preparation

Chemical cross-linking with DSSO (Kao, AH, et al., 2011; Mendes, ML, et al., 2019) followed by mass spectrometry was used to examine CheA domain arrangements in the free kinase and ternary complex (Tar receptor foldon:CheA:CheW). For the sample of CheA alone, ∼50 µg of of the protein was cross-linked in solution (50 mM HEPES pH 7.5, 250 mM KCl, 10% glycerol, 100 µM ADP, 5 mM MgCl_2_) and then dried. The sample was then denatured and reduced with the addition of 20 µl 6M Guanidinium-HCL, 50 mM Ammonium bicarbonate and 2 µl 0.110M DTT (final concentration = 10 mM DTT) and incubated at 60°C for 1 hour. The sample was then alkylated with the addition of 2.5 µl of 0.55 M IDA followed by 45 minutes of incubation at room temperature in the dark. The alkylation reaction was quenched with 5.5 µl 0.2 M DTT. The sample was then diluted by the addition of 100 µl of 50 mM ammonium bicarbonate. 12.5 µl of Trypsin at 0.2 µg/µl (2.5 µg) was added and incubated overnight at 37°C. The sample was heated to 90°C for 5 minutes to deactivate the trypsin. A second digestion with chymotrypsin was then carried out by the addition of 5 µl of 1 µg/µl chymotrypsin followed by an overnight incubation at 37°C. The digestion was stopped with the addition of 150 µl of 0.5% TFA. The pH after addition was < 2.5. The samples were then applied to a SepPak C18 1cc/ 50 mg (waters) cartridge for cleanup.

For the ternary complex, a protein sample containing a 1:1:1 ratio of the composite proteins was incubated and cross-linked in solution (50 mM HEPES pH 7.5, 250 mM KCl, 10% glycerol, 100 µM ADP, 5 mM MgCl_2_) and then run on an SDS page gel. A gel band containing ∼50 µg the ternary complex was excised from the gel for MS/MS analysis. First, the gel band was washed with 400 µl ddH_2_O, then 400 µl of 50% ACN and 50 mM Ambic, then 400 µl 100% CAN. The gel was exposed to air in a chemical hood until dryness. The sample was then reduced with the addition of 200 µl of 10 mM DTT in 100 mM Ambic at 60°C for 1 hour. The sample was alkylated with the addition of 200 µl of 55 mM IDA in 100 mM Ambic and incubated for 45 minutes at room temperature in the dark. The washing steps were then repeated. 120 µl Trypsin at 10 ng/µl (1.2 µg) in 50 mM Ambic and 10% ACN was added and then incubated on ice for 20 minutes. The sample was then overlayed with another 150 µl of 50 mM Ambic with 10% ACN and incubated overnight at 37°C. After 18 hours of digestion, the reaction was stopped with the addition of 5 µl of 100% Formic acid. The sample was then extracted from the gel by first vortexing in a mixture of 200 µl of 50% ACN and 5% formic acid for 30 minutes at 1800 rpm followed by sonication for 10 minutes. This step was repeated once more and the sample was dried in a speed vacuum. The sample was then reconstituted with 45 µl of 50 mM Ambic and heated to 90°C for 5 minutes. The second digestion with chymotrypsin was then conducted by adding 5 µl chymotrypsin at 1 µg/µl (3 µg) and incubated overnight at 37°C. The digestion was stopped with the addition of 2 µl of 100% Formic acid. The resulting pH was <2.5. The sample was dried to dryness in a speed vacuum and then resuspended into 0.5% Formic acid.

#### NanoLC-MS/MS analysis

Peptides and cross-linked peptides were analyzed using an UltiMate3000 RSLCnano (Dionex, Sunnyvale, CA) coupled to an Orbitrap Fusion (Thermo-Fisher Scientific, San Jose, CA) mass spectrometer equipped with a nanospray Flex Ion Source. Each sample was loaded onto an Acclaim PepMap 100 C_18_ trap column (5 µm, 100 µm × 20 mm, 100 Å, Thermo Fisher Scientific) at 20 *μ*L/min of 0.5% formic acid (FA). After 3 minutes, the valve switched to allow peptides to be separated on an Acclaim PepMap C18 nano column (3 µm, 75µm x 25cm, Thermo Fisher Scientific), in a 120 min gradient of 5% to 40% B at 300 nL/min. The Orbitrap Fusion was operated in positive ion mode with nano spray voltage set at 1.7 kV and source temperature at 275 °C. External calibrations for Fourier transform, ion-trap and quadrupole mass analyzers were performed prior to the analysis.

Samples were analyzed using the CID–MS2–MS3 workflow, in which peptides with charge states 4–10 were selected for CID MS2 acquisitions in Orbitrap analyzer with a resolution of 30,000 and an AGC target of 5 × 10^4^. MS scan range was set to 375–1575 m/z and the resolution was set to 60,000. The precursor isolation width was 1.6 m/z and the maximum injection time was 100 ms. The CID MS2 normalized collision energy was set to 25%. Targeted mass-difference-dependent CID–MS3 spectra were triggered for acquisition in the ion trap with CID collision energy of 35%; AGC target of 2 × 10^4^ when a unique mass difference (Δ=31.9721 Da) was observed in the CID–MS2 spectrum. MS2 isolation window of 3 m/z with the maximum injection time set to 100 ms. All data were acquired under Xcalibur 3.0 operation software and Orbitrap Fusion Tune Application v. 2.1 (Thermo-Fisher Scientific).

#### Data processing, protein identification and data analysis

All MS, MS2 and MS3 raw spectra from each sample were searched using Proteome Discoverer 2.2 (Thermo-Fisher Scientific, San Jose, CA) with XlinkX v2.0 algorithm for identification of cross-linked peptides. The search parameters were as follow: three missed cleavage for full trypsin digestion or double digestion with fixed carbamidomethyl modification of cysteine, variable modifications of methionine oxidation. The peptide mass tolerance was 10 ppm, and MS2 and MS3 fragment mass tolerance was 20 ppm and 0.6 Da, respectively. The *Ecoli* database with added targeted protein sequences was used for PD 2.2 database search with 1% FDR for report of cross-link results. In addition, the search was also performed using the full sequences of the proteins including the recombinant tags. Identified cross-linked peptides were filtered for Max. XlinkX Score >20 containing at least two identified MS3 spectra for each pair of cross-linked peptides. Results of the search with were exported by the software as a spreadsheet.

### Pulse Dipolar ESR Spectroscopy (PDS)

PDS measurements were conducted by four pulse double electron electron resonance (DEER) at 60 K based on a 17.3 GHz FT ESR spectrometer (Borbat, PP, et al., 1997) modified for PDS ESR (Borbat, PP, et al., 2013) and at 34 GHz in Q-band on on a Elexsys E580 spectrometer equipped with a 10W solid state amplifier (150W equivalent TWTA) and Arbitrary Waveform Generator (AWG) to generate MW pulses at the detection and pumping frequency offsets to suppress spurious echoes. Q-band DEER measurements were performed at 60 K (ER 4118HV-CF10-L FlexLine Cryogen-Free VT System) in an EN 5107D2 cavity using four pulses (*π/2*-τ_1_-*π*- τ_1_-*π*_pump_- τ_2_-*π* - τ_2_-echo) with 16-step phase cycling. Data were collected on 100 µM protein over 16 hours for each sample. The signal background was approximated by a polynomial function in the semi-log scale and subtracted out (Borbat, PP, et al., 2002). Noise from the time domain data was then removed by the WavPDS method (Srivastava, M, et al., 2017) (a wavelet denoising procedure for PDS) and distance distributions of spin separations calculated by the new singular-value decomposition (SVD) method (Srivastava, M, et al., 2017; Srivastava, M, et al., 2019) developed to solve ill-posed problems such as for the PDS signal and estimate uncertainty in their measurement.

### Model Building of the Ternary Complex

#### Rosetta Loop modeling

For the dipped CheA models, ten and seven residues were selected to be modeled for reconnecting the P3-P4 (L3) loop and the P4-P5 (L4) loop, respectively. More residues were used in loop modeling based on further distance between the domains of CheA. The chain breaks were located one residue further away from P4 domain in both loops of both CheA models to allow more dynamic motion of the domain in this study. In L3 loop, the shorter range of residues to be modeled also resulted in less disruption to the dimerization characteristic of the P3 domain. In productive run, 250 KIC build attempts were made with 500 models generated. The best model was selected based on the Rosetta Energy Unit (REU) score and finalized after structural comparison with the other top models. The new P3-P4 and P4-P5 loop conformations of the dipped CheA models were reintroduced in the original models before Rosetta Relax.

#### Rosetta Relax and RosettaEPR

In Rosetta Relax productive run of the dipped CheA-complex (CheW and receptors) model was set to generate 100 models with five cycles of minimization and sidechain repack. Furthermore, in the dipped CheA-complex model, SDSL-PDS distance restraints (Table 1) were included in the relax protocol. A suitable function based on the “motion-on-a-cone” model developed by Alexander et al. (Alexander, N, et al., 2008) was implemented in RosettaEPR or the PDS restraints The function describes the relationship between the experimental measured spin label distance (dSL) and Cβ-Cβ distance (dCβ) in which an acceptable range of dCβ is determined from dSL (dCβ ϵ [dSL-12.5 Å, dSL +2.5Å]). Top scoring models based on the REU scores were selected and analyzed. The best model of the dipped CheA complex has an REU score of −4845.83 a, with the average score out of 100 models at −4487.6. Top ten scoring models of dipped CheA were selected for analysis. In comparison to P3, P4 and P5 domains of the best model, the rest of the models have a root mean square deviation (RMSD) of 1.90-2.75 Å (chain A) and 2.09-4.63 Å (chain B) for 373 Cα atoms.

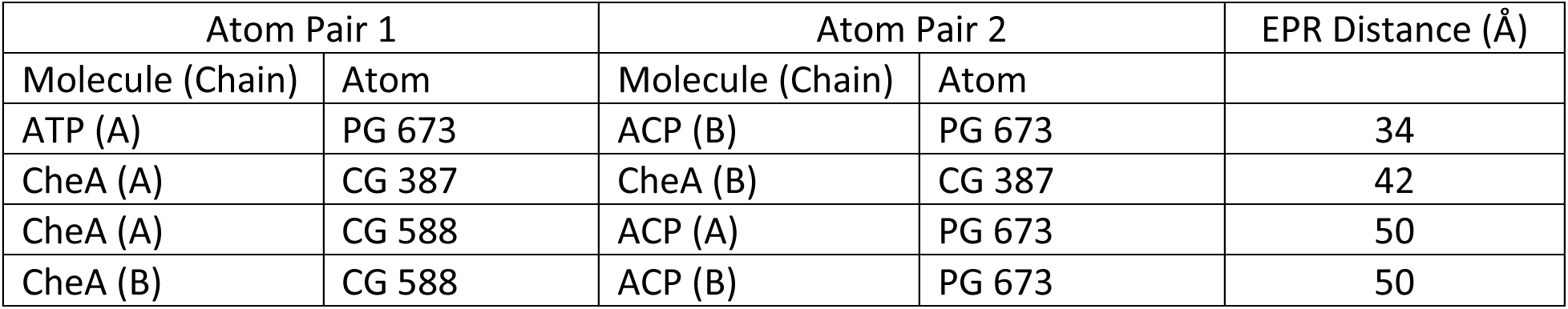

| Atom Pair 1 |  | Atom Pair 2 |  | EPR Distance (Å) |
| --- | --- | --- | --- | --- |
| Molecule (Chain) | Atom | Molecule (Chain) | Atom |  |
| ATP (A) | PG 673 | ACP (B) | PG 673 | 34 |
| CheA (A) | CG 387 | CheA (B) | CG 387 | 42 |
| CheA (A) | CG 588 | ACP (A) | PG 673 | 50 |
| CheA (B) | CG 588 | ACP (B) | PG 673 | 50 |

#### Modeling of P1 and P2 in the inhibited complex

As supported by PDS data, the crystallographic P1 dimer was placed in the Rosetta model with CHIMERA in order to be consistent with the SAXS envelope and cross-linking data. The P2 domains were subsequently added symmetrically under similar considerations. The L1 and L2 loops were built with EOM (Bernado, P, et al., 2007) as C*α* traces. Families of configurations were evaluated against the SAXS data with FoXS (Schneidman-Duhovny, D, et al., 2016)after replacing the traces with the correctly sequenced polypeptide. Six general families of loop positions that extended into different regions of the SAXS envelopes were considered in this manner. Only those configurations that extended the L1 loops below the P1 domains sufficiently extended the radius of gyration to produce reasonable agreement with the SAXS data. With the L1 and L2 loops explicitly built and the receptor foldons incorporated, Rosetta Relax was used to produce 100 new models under RosettaEPR constraints of the PDS distances and bounded-function constraints of the cross-linking data. Protein cross-links were assumed to occur over a range of 20 ± 7 Å. Each of the 100 final models was evaluated against the SAXS data with FoXs. The model with the lowest Rosetta energy and the model with the closest agreement to the SAXS data have a REU score of −5863.93 and −5679.7 respectively. Their average scores out of 100 models is at −5333.27. Disagreeable conformations and with high (> 0) Rosetta energy scores were excluded. The best model that maintained the expected domain interfaces was then subjected configurational sampling about the L4 linker, while keeping P5, CheW and receptor foldons as rigid bodies. This sampling, implemented in MultiFoxS (Schneidman-Duhovny, D, et al., 2016) maximized agreement with the SAXS data and the subsequent model was fit against the SAXS-derived molecular envelope in Chimera. Restraint distances in the final model are given in Figure 8 – figure supplement 2.

## Acknowledgements

This work was supported by grants from the National Institutes of Health: R35GM122535 to BRC and P41GM103521 for the National Biomedical Center for Advanced ESR Technologies (ACERT) to JHF, and was supported by the European Union under a Marie-Sklodowska-Curie COFUND LEaDing fellowship to ARM and a National Science Foundation Graduate Research Fellowship Program (NSF-GRFP) grant to ARM (2014155468) We thank Estella Yee for help with kinetics fits, the Cornell High Energy Synchrotron Source (CHESS) for access to data collection facilities. CHESS is supported by NSF award DMR-1332208 and NIH/NIGMS award P30GM103485.

## Author contributions

A.R.M., T.K., W.Y., Z.M., A.B., and B.R.C. designed research; A.R.M., T.K., M.S., W.Y., Z.M., P.P.B., J.C., S.Z., and B.R.C. conducted research; A.R.M., T.K., M.S., W.Y., Z.M., P.P.B., J.C., S.Z., J.H.F, A.B., and B.R.C. analyzed data; A.R.M, T.K., and B.R.C. wrote the manuscript with input from all authors.

## Supplementary Figure Legends

**Figure 1 – figure supplement 1.**
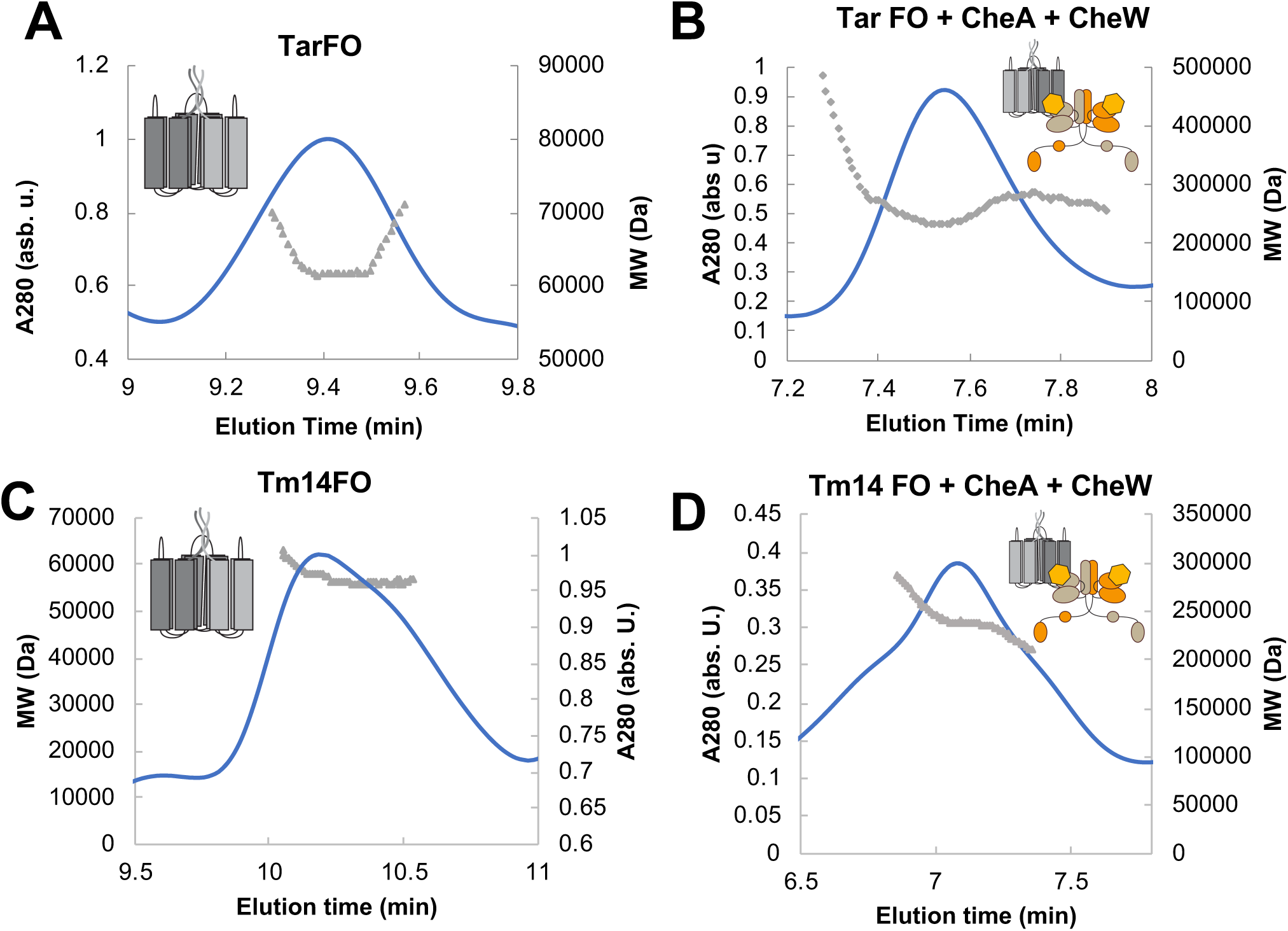
Preliminary purification and characterization of receptor foldons and their complexes. (A) MALS experiments indicate that the Tar foldon has a MW (60 kDa) that matches expectations for a trimer and produces higher molecular-weight complexes with *Thermotoga maritima* (*Tm*) CheA and CheW. (C) MALS experiments indicate that the Tm14 foldon has a MW (60 kDa) that also corresponds to a trimer-of-single-chain-dimers and (D) produces higher molecular-weight complexes with Tm CheA and CheW.

**Figure 1 – figure supplement 2.**
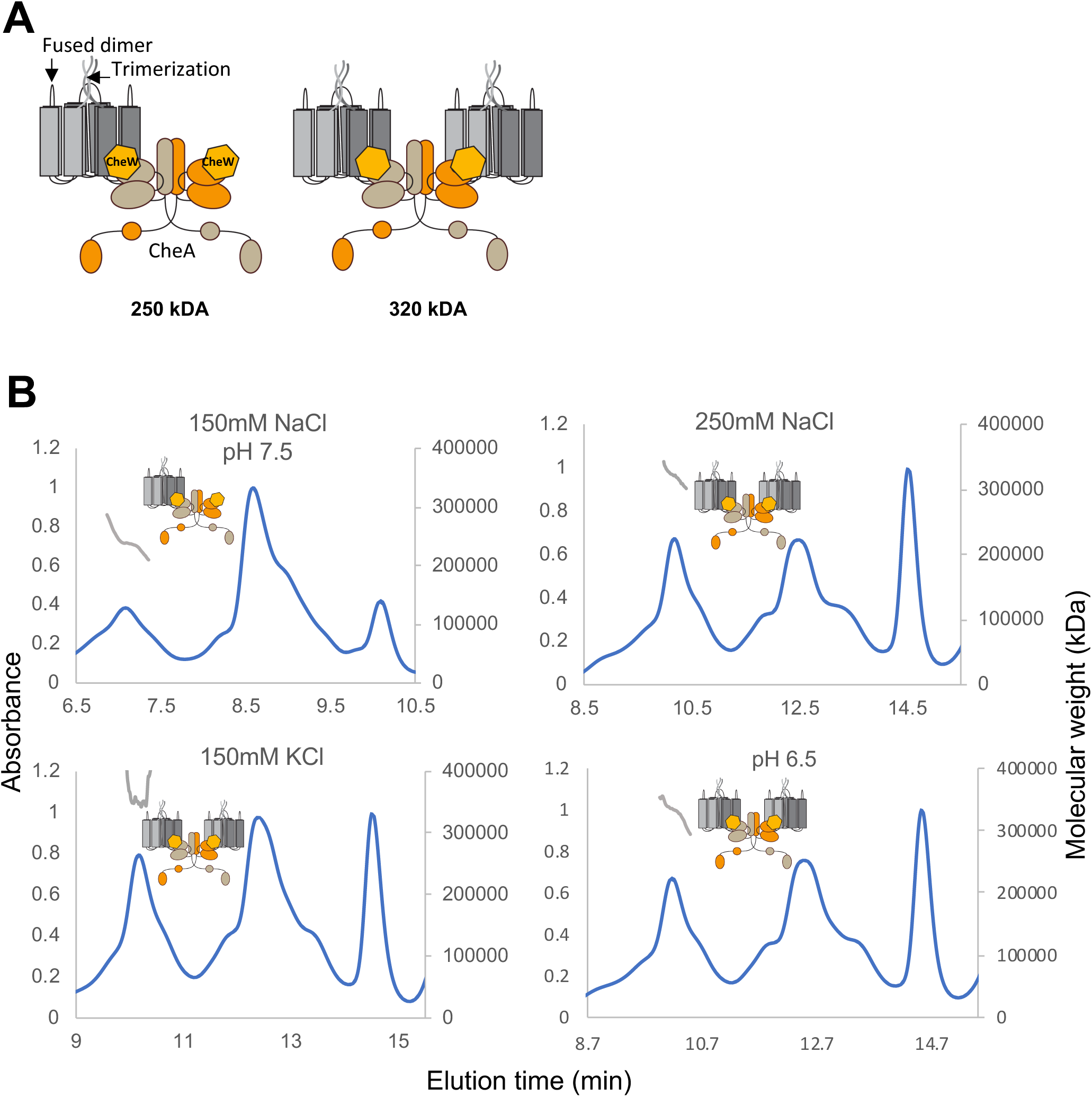
MALS experiments demonstrate that specific buffer conditions encourage interactions among the Tar foldons and CheA/W. (A) The foldons primarily form two particles in solution: MW = 250 kDa or MW = 320 kDa, the latter consistent with the size of a core signaling unit. (B) Standard buffer conditions produce ternary complexes that are primarily 250 kDa. Increasing NaCl concentration, replacing NaCl with KCl, or decreasing pH favors the production of the 320 kDa species.

**Figure 1 – figure supplement 3.**
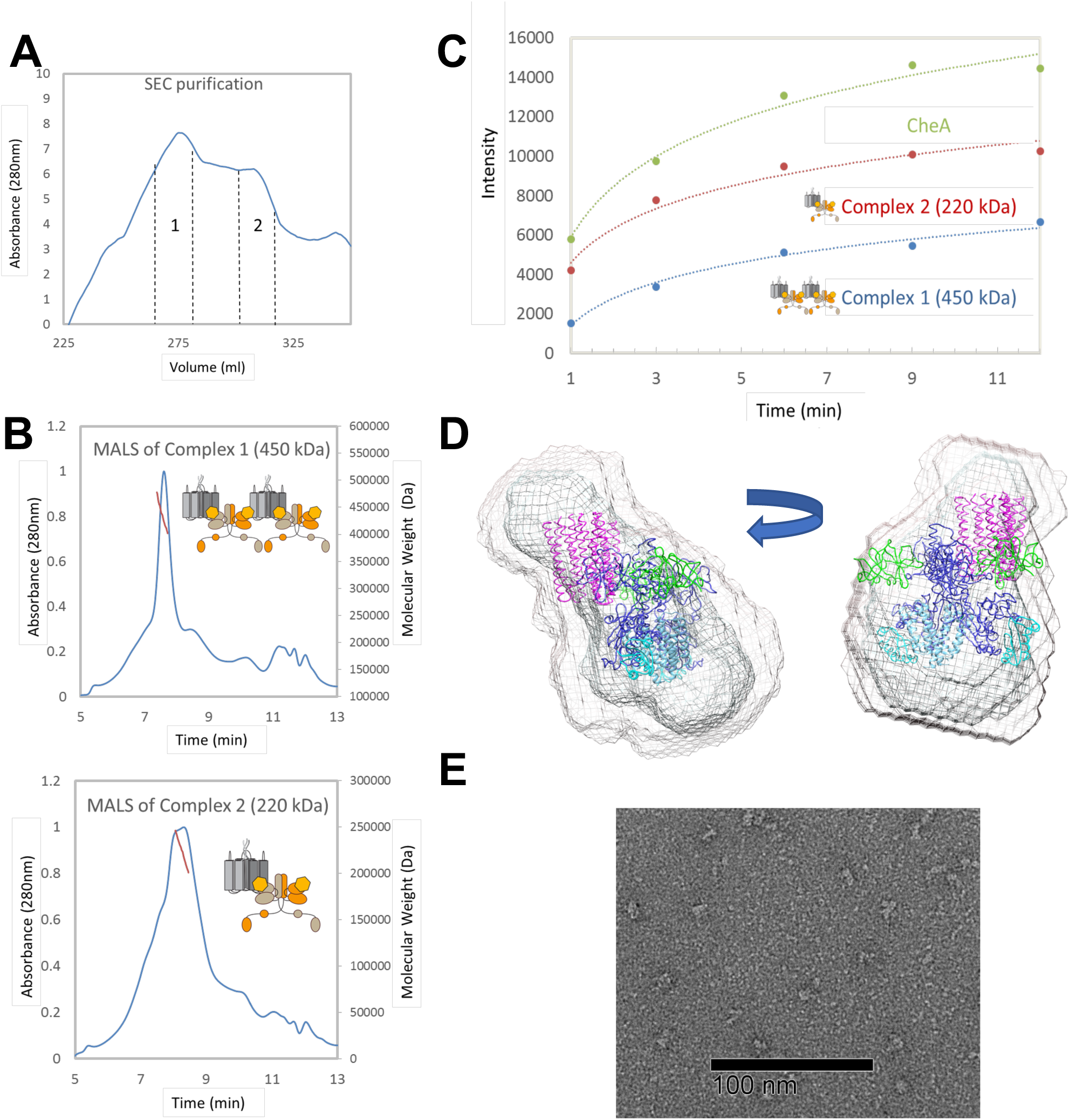
Analyses of ternary complexes consisting of the Tm14 foldon with *Tm* CheA and *Tm* CheW. (A) Top: Size-exclusion chromatography of a mixture of Tm14 foldon with CheA and CheW produces high-molecular weight peaks (denoted 1 and 2). Quantitative SDS-PAGE analysis of peak 1 indicates Tm14 foldon:CheA:CheW in a roughly equal subunit ratio. (B) MALS experiments with complexes in Peak 1 denote particles of MW that corresponds to four CheA subunits, 4 CheW subunits and two receptor foldons. MALS of Peak 2 indicates that the particles consist of one CheA dimer with two CheW subunits and one receptor foldon. (C) Complexes from Peak 1 and Peak 2 deactivate *Tm* CheA in ^32^P-ATP phosphorylation. The 450 kDa complex deactivates CheA ∼3 fold and the 220 kDa complex deactivates CheA ∼1.5 fold; CheA subunit concentration was kept constant. (D) SEC-SAXS of the complexes in Peak 1 produces a molecular envelope that fits the expected size of the predicted components. Model shown is an approximate assembly based on crystal structures and array models to gauge spatial extent. (D) Negative-stain electron microscopy reveals particles in Peak 1 are relatively monodispersed and around 100 Å in diameter; however, the particles were too heterogenous for 2D classification.

**Figure 1 – figure supplement 4.**
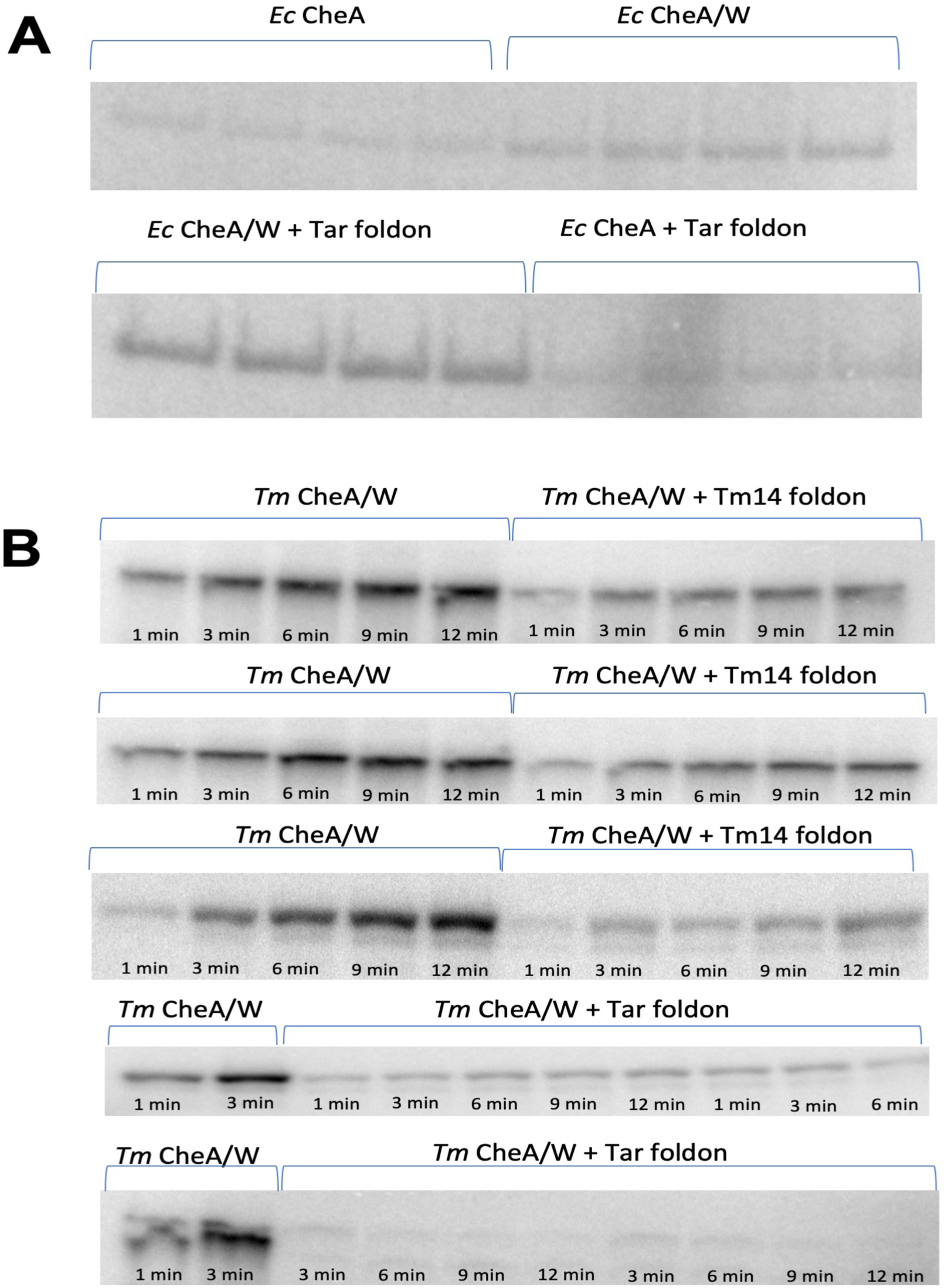
Autoradiographs of the CheA autophosphorylation experiments from Figure 1. (A) Radioisotope assays with Ec CheA show CheW-dependent activation by the Tar foldon. Assays performed in parallel and imaged together for comparison of CheW-dependent activation of free Ec CheA and in signaling units. (B) Radioisotope assays with Tm CheA demonstrate that the enzyme is deactivated by both foldons, but almost completely deactivated by the Tar foldon.

**Figure 1 – figure supplement 5.**
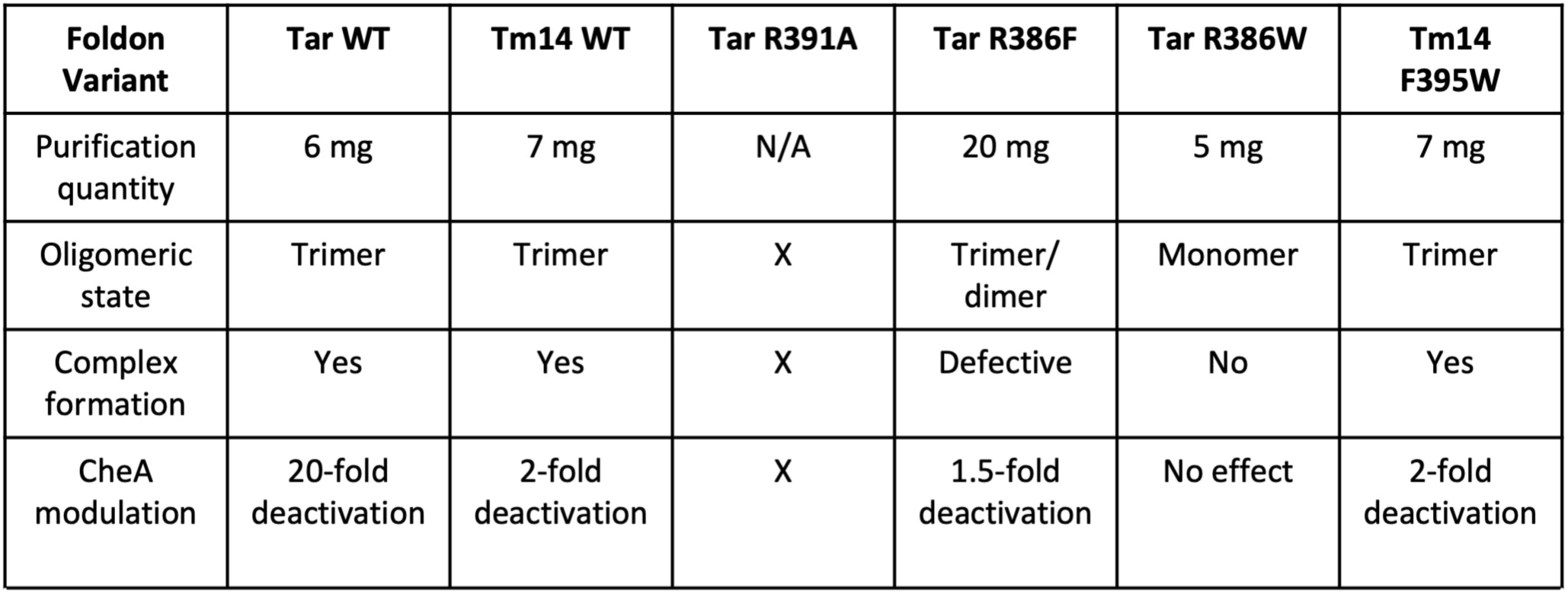
Properties of single-residue foldon variants and their complexes with CheA.

**Figure 2 – figure supplement 1.**
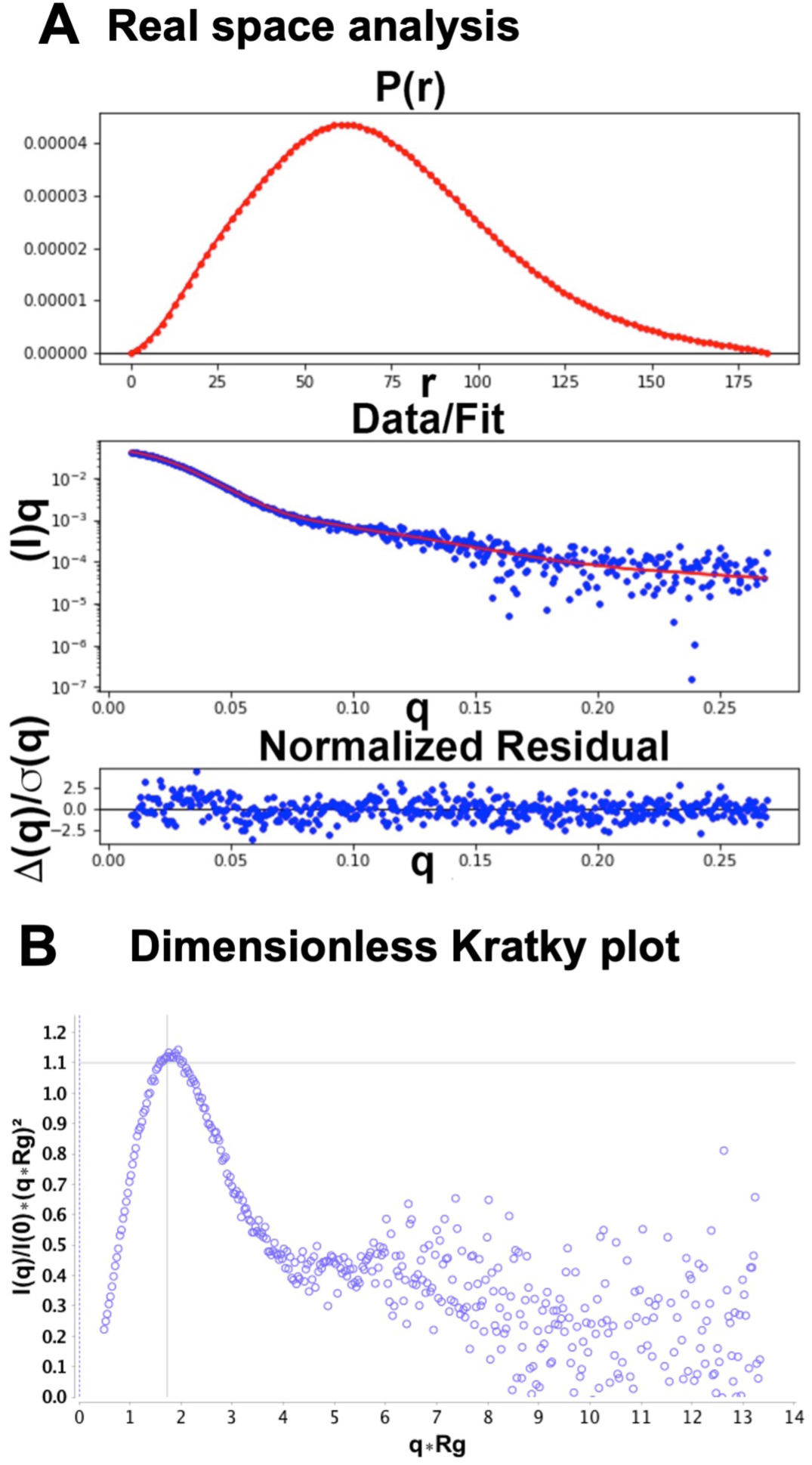
SEC-SAXS data of the cross-linked foldon complex (A) Real-space analysis of the particle by GNOM after SVD decomposition of the SEC-SAXS profile (Volume of Correlation MW = 305 kDa; Porod volume MW = 328 kDa; DAMAVER MW = 339 kDa, X2 = 1.34 Dmax = 201 Å Radius of Gyration (Rg) = 55.4 Å, Normalized Spatial Discrepancy (NSD) = 1.34; Gunier Rg = 55.4 Å; GNOM Rg = 53.8 Å). (B) Dimensionless Kratky plot of the cross-linked complex from ScÅtter indicate a globular shape of the particle. Globular particles show an intersection of the x and y variables at the cross-hairs of the plot.

**Figure 3 – figure supplement 1.**
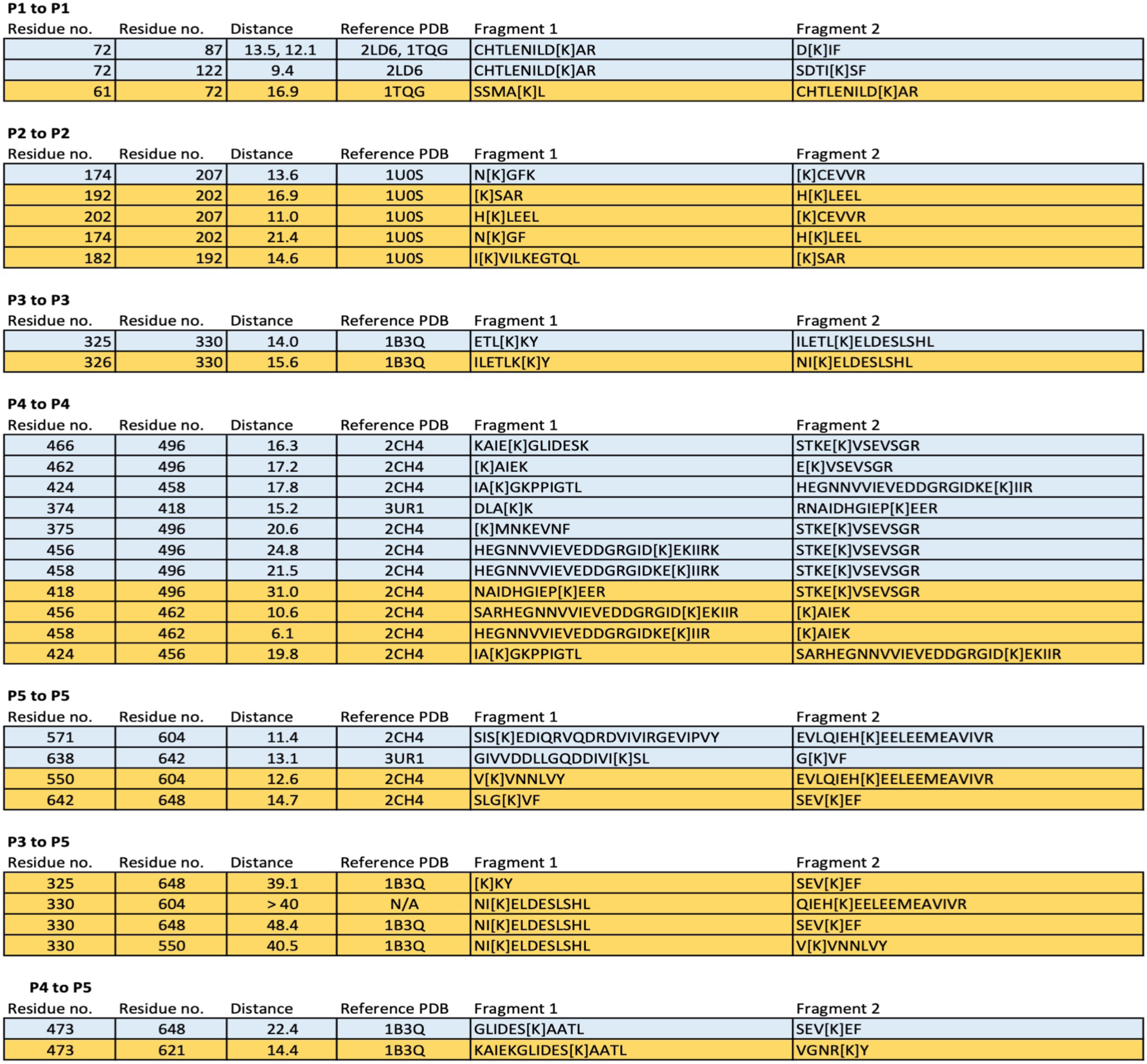
Inter-domain cross-links in free CheA. Cross-linked peptides in yellow are found in the free kinase, but not the foldon complex.

**Figure 3 – figure supplement 2.**
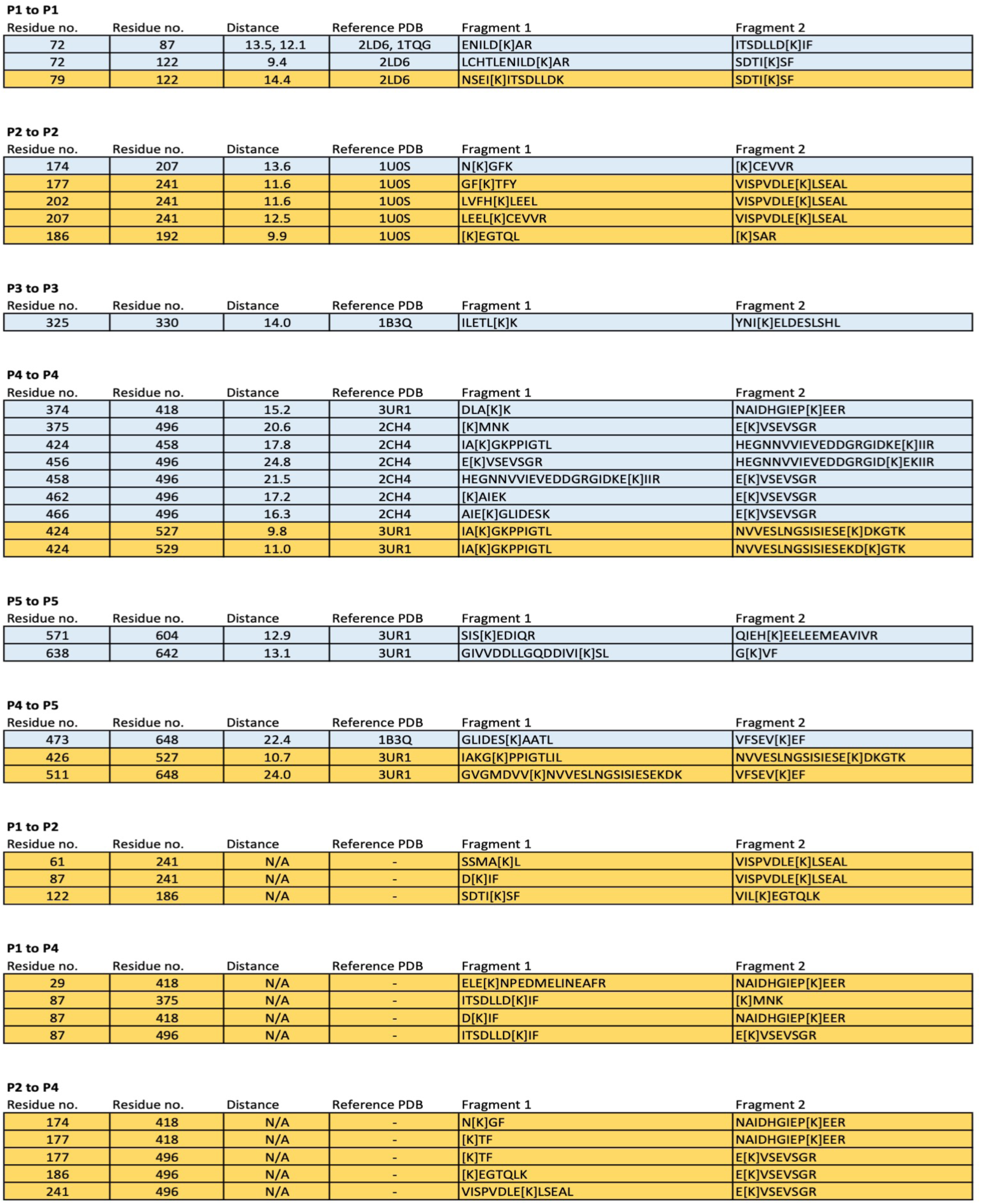
Inter-domain cross-links in the receptor foldon complex. Cross-linked peptides in yellow are found in the foldon complexes, but not the free kinase.

**Figure 7 – figure supplement 1.**
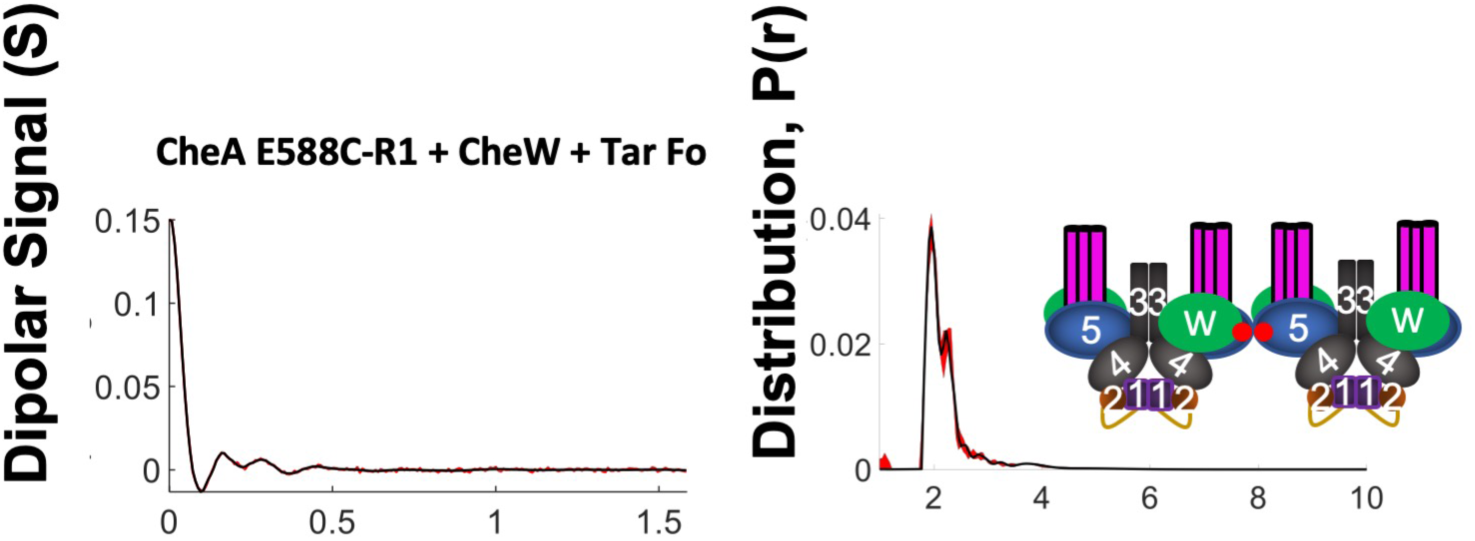
DEER spectroscopy experiments of *Tm* CheA E588C-R1 with CheW and Tar foldon. A rapid echo decay and strong oscillation indicates a short distance in the time domain data, as resolved at right in the distance distribution. CheA is known to associate through its P5 domains in a manner that would bring opposing 588C-R1 sites in close proximity (shown by red dots in the schematic). In absence of other spin sites, the close 588C-R1 positions dominate the PDS signal.

**Figure 8 – figure supplement 1.**
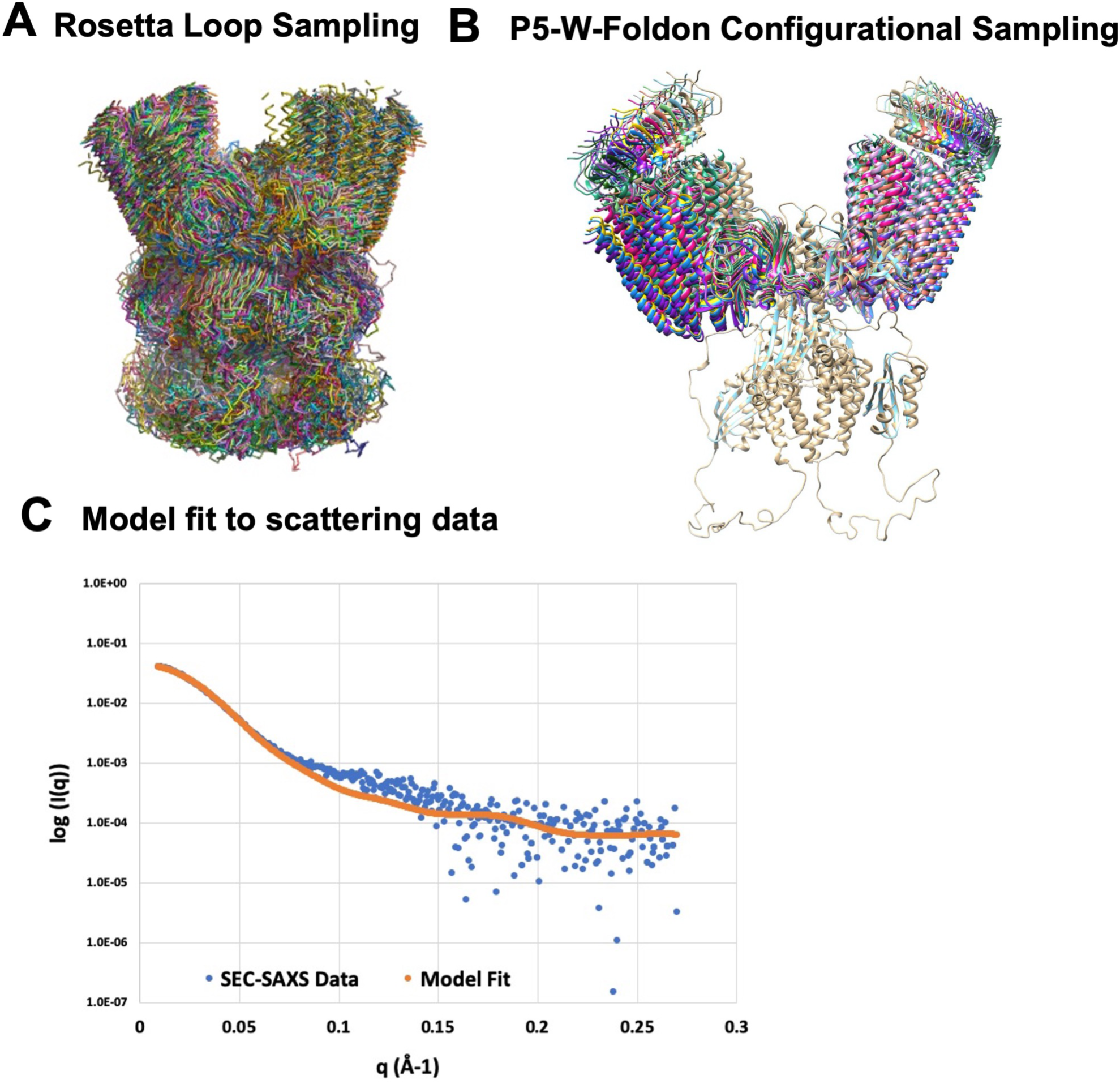
Modeling of the foldon complex. (A) Overlay of C*α* traces representing conformations sampled in Rosetta under constraints from PDS and cross-linking. Conformations were evaluated for their agreement with SAXS data and low Rosetta energy scores. (B) The highest scoring model in (A) was subjected to loop perturbation about the P4-P5 hinge (L4), while treating P5-CheW-receptor foldon units as rigid units on each CheA subunit, and then subsequently evaluated against the SAXS data. Highest scoring model (X^2^ = 4.5) fit to the scattering data is shown in (C).

**Figure 8 – figure supplement 2.**
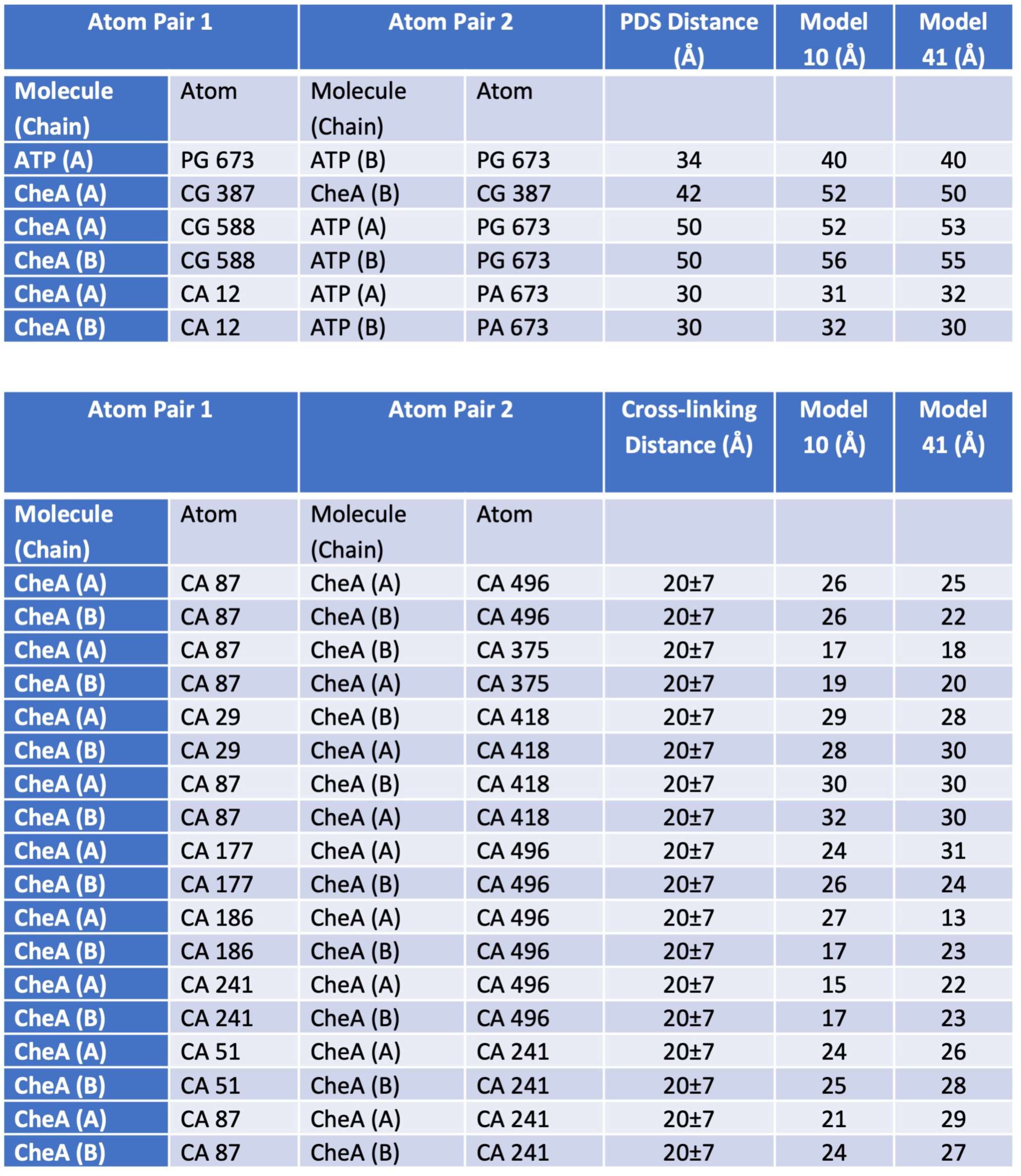
Summary of model restraints in Rosetta refinements based on PDS measured distances (top) and unique DSSO cross-links formed in the foldon-receptor complexes (bottom). Model 10 represents the model with the lowest Rosetta energy (−5863 Rosetta Energy Units (REUs)); model 41 represents the model with the closest agreement to the SAXS data (−5680 REU).

